# Synthetic cannabinoid receptor agonists inhibit the cardiac voltage-gated potassium channel hERG

**DOI:** 10.64898/2025.12.03.692044

**Authors:** Nina E. Ottosson, Damon J. A. Frampton, Tanadet Pipatpolkai, Caitlyn Norman, Urban Karlsson, Amaia Jauregi-Miguel, Maryke Venter, Akshay Sridhar, H. Peter Larsson, Henrik Gréen, Sara I. Liin

## Abstract

Synthetic cannabinoid receptor agonists (SCRAs) is a large group of structurally diverse designer drugs (analogues of controlled substances) associated with intense and sometimes fatal intoxication. Cardiac symptoms including tachycardia and arrhythmia are common consequences of SCRA consumption. However, little is known about the mechanisms through which SCRAs may perturb cardiac rhythm. Here we used electrophysiological techniques to screen 36 SCRAs on two ion channels responsible for cardiomyocyte repolarization, hERG (also called K_V_11.1) and K_V_7.1/KCNE1. We report that the majority of tested SCRAs inhibited hERG, primarily by reducing channel conductance, and some also inhibited K_V_7.1/KCNE1. *In silico* data suggests that SCRAs use a known drug binding site in the pore of the hERG channel, shared by established hERG blockers like astemizole, where a planar SCRA molecule lays perpendicular to the ion conducting pathway. Experimental and *in silico* data identify SCRA structural features associated with prominent inhibitory effects on hERG, with chemical moieties allowing bond formation and/or the ability to fit into the vestibule being important. Structure-activity-relationships for SCRA effects on hERG, K_V_7.1/KCNE1 and the cannabinoid receptor 1 (CB_1_) varied, demonstrating the importance of assessing SCRA effects on multiple potential targets. In conclusion, we found SCRAs to be inhibitors of cardiac voltage-gated potassium channels important for cardiomyocyte repolarization. These results offer mechanistic insight into potentially detrimental SCRA effects on the heart and highlight the urgency of more extensive investigation of SCRAs on cardiac function.

## Introduction

Synthetic cannabinoid receptor agonists (SCRAs, exemplified by JWH-018 in Fig. 1A), commonly known as “spice” or “K2”, is a large class of new psychoactive substances (NPS) designed to mimic the effects of Δ^9^-tetrahydrocannabinol (Δ^9^-THC) (Fig. 1B), the main psychoactive component of cannabis ^1^. The SCRAs available on the recreational drug market is constantly changing with new compounds emerging each year to circumvent national and international legislation, posing international threats and challenges to the communities. For newly emerging SCRAs, chemical modifications may be seen at the core, tail, head (also known as linked group), or linker of the SCRA molecule (Fig. 1C). As of 2025, 277 different SCRAs have been reported to the European Union early warning system, and it is the largest group of NPS being monitored by the European Union Drug Agency (EUDA) ^2^.

**Figure 1:**
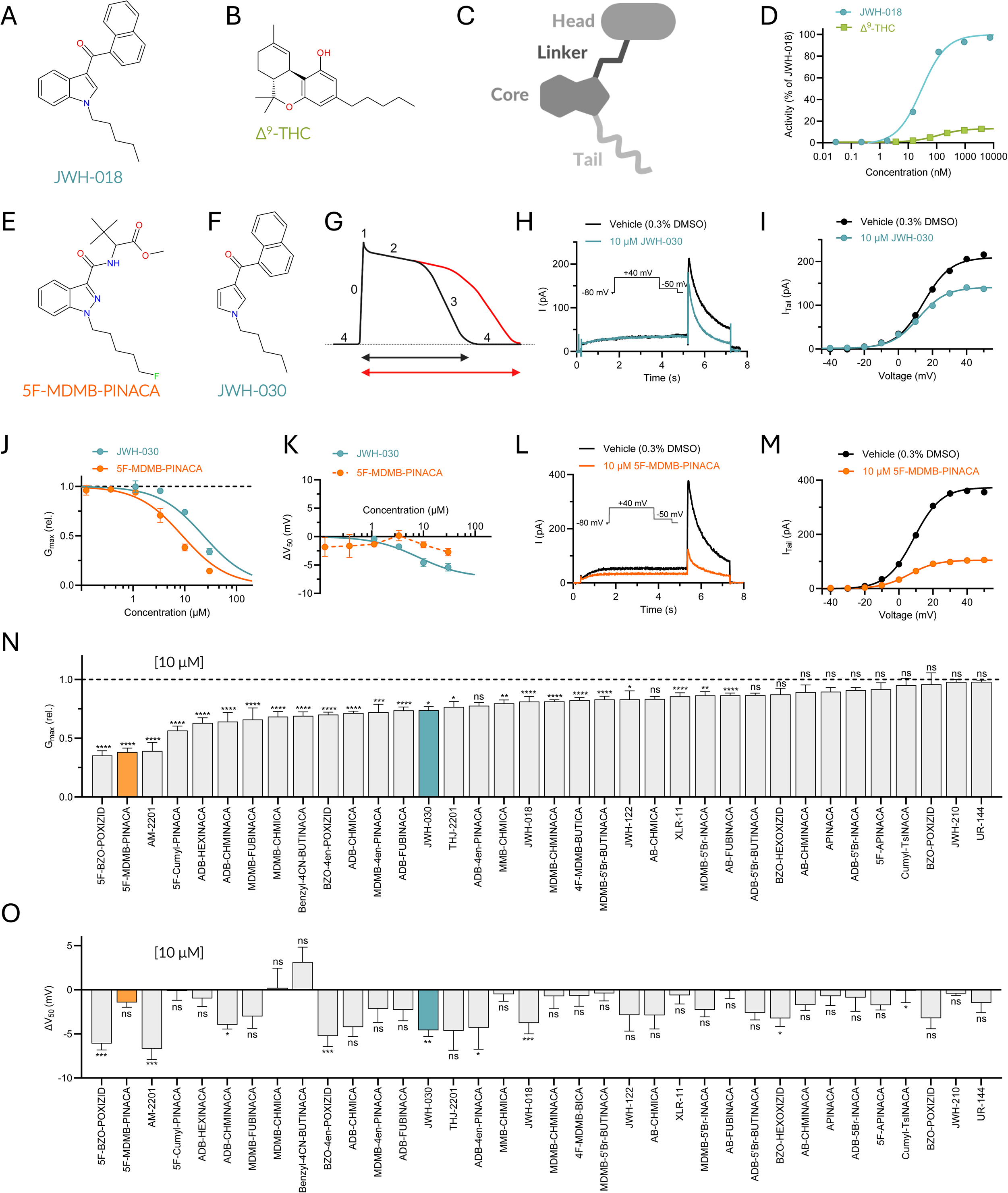
The majority of tested SCRAs inhibit hERG. (A-B) Chemical structure of the SCRA JWH-018 (A) and Δ^9^-THC (B). **(C)** Schematic illustration of the division of a generic SCRA into four key building blocks: the head, linker, core, and tail. **(D)** CB_1_ receptor activation of JWH-018 and Δ^9^-THC, where data is normalized to the maximal response of JWH-018 (used as reference), arbitrarily set at 100%. **(E-F)** Chemical structures of the SCRAs 5F-MDMB-PINACA (E) and JWH-030 (F). **(G)** Schematic illustration of an action potential from a ventricular cardiomyocyte, with its five phases (0-4) indicated, where phase 3 represents the repolarization phase. The solid black line represents an action potential under normal conditions, the solid red line represents action potential prolongation due to compromised repolarization (e.g, reduced I_Kr_ or I_Ks_). **(H-I)** Representative effect of JWH-030 on hERG using the pulse protocol shown as insert (H), with corresponding G(V) curve (I). **(J-K)** Concentration-response relationship for G_max_ (rel) (J) or ΔV_50_ (K) effect of JWH-030 and 5F-MDMB-PINACA. n = 6-8 for JWH-030; n = 4-17 for 5F-MDMB-PINACA. **(L-M)** Same as in H-I but for 5F-MDMB-PINACA. **(N-O)** Average effect of 10 µM of the indicated SCRAs on G_max_ (N) and V_50_ of hERG (O). n = 3-13. Statistics denote adjusted p-values for comparisons to vehicle: p < 0.05 (*), p < 0.01 (**), p < 0.001 (***), p < 0.0001 (****). All average data are shown as mean ± SEM. Small error bars are covered by symbols. Best fits as in Table II.

SCRAs have been estimated to be used by 0.2-4% of the population ^3^ with much higher use reported in prisons ^4^, rough-sleeping populations ^5^, and populations with regular drug testing, including the military ^6^ and people on probation/parole ^7^. SCRAs are often smoked as infused herbal material or vaped as infused e-cigarette oils ^8^. There have also been many reports of the addition of SCRAs to cannabis products on the market ^9,10^ and SCRAs found in drug samples sold as opioids or benzodiazepines ^11^, meaning people who use SCRAs are often unaware of the actual SCRAs they are using or may not even know they are using SCRAs.

The high variability of the market and lack of pharmacological data of newly emerging drugs significantly increases the risk of harms. SCRAs are typically full agonists of the cannabinoid 1 (CB_1_) receptor, the receptor responsible for the psychoactivity of cannabis, whereas Δ^9^-THC is only a partial agonist ^1^. SCRAs are often significantly more potent on the CB_1_ receptor than Δ^9^-THC (exemplified by JWH-018 in Fig. 1D) and have many adverse effects reminiscent of psychostimulants rather than cannabis, including psychosis, seizures, and cardiac arrhythmia ^12^. Cardiovascular-related complications of SCRAs are the most common symptoms presented at accident and emergency departments ^13,14^ and common causes of death ^15–17^. For instance, 5F-MDMB-PINACA (Fig. 1E, also known as 5F-ADB), has been implicated in several fatal intoxications ^18–20^ and linked to cardiac symptoms ^21^. Thus, there is a substantial body of evidence linking SCRA intoxication to potentially fatal cardiac arrhythmias ^22^, with QT prolongation on the electrocardiogram (ECG) reported in several studies e.g., ^23,24,25^.

Despite frequent reports associating SCRA intoxication with arrhythmias, little is known about how SCRAs cause these adverse effects on a mechanistic level ^22^. As cardiac arrhythmias are electrophysiological disorders, cardiac ion channels may be implicated. As the first example supporting this, an electrophysiological study reported that JWH-030 (Fig. 1F) inhibits the voltage-gated potassium channel hERG (K_V_11.1) and prolongs the QT interval in rats ^26^. Moreover, a recent integrative computational and electrophysiological study show that 5F-APINACA (also called 5F-AKB48) and ADB-FUBIATA inhibit hERG and find a side pocket lateral to the central cavity for these two SCRAs ^27^. Another study used machine learning models to propose hERG binding of several SCRAs, though no electrophysiological experiments were conducted to confirm the predictions ^28^. These three studies provide the first indication on the interaction between SCRAs and hERG, highlighting the importance of more extensive studies.

The hERG channel, together with the K_V_7.1/KCNE1 channel, is responsible for phase 3 repolarization of the action potential in ventricular cardiomyocytes: hERG carrying the rapidly activating delayed rectifier current I_Kr_, whereas K_V_7.1/KCNE1 carries the slowly activating delayed rectifier current I_Ks_ (Fig. 1G). Impaired function of hERG or K_V_7.1/KCNE1 can lead to a prolonged action potential duration (indicated in red in Fig. 1G), early afterdepolarizations and arrhythmic beats ^29,30^. This is observed in ventricular arrhythmias such as Torsades de Pointes, a life-threatening polymorphic tachycardia that is a consequence of both drug-induced Long QT Syndrome (LQTS) following hERG channel blockade and as congenital LQTS caused by deleterious mutations in the genes encoding hERG and K_V_7.1/KCNE1 ^29,30^. However, whether SCRAs other than JWH-030, 5F-APINACA, and ADB-FUBIATA inhibit hERG remains unclear. In addition, the underlying mechanisms of action for most SCRAs and their putative compounding proarrhythmic effects on K_V_7.1/KCNE1 are unknown. This lack of understanding limits risk stratification of SCRAs, medical treatment protocols, and harm-reducing actions, such as implementing legislative controls. Thus, our study aims to address this gap in knowledge by determining the effects and mechanisms of a broad set of SCRAs on hERG and K_V_7.1/KCNE1 channels.

## Results

### SCRAs inhibit the hERG channel by reducing channel conductance

To assess the effect of a large set of SCRAs on hERG (see SI Table I and SI Figure S1 for a list of the 36 included SCRAs and corresponding structures), we used the automated planar patch-clamp (APC) technique on mammalian Chinese hamster ovary (CHO) cells stably expressing human hERG. All 36 SCRAs were evaluated in concentration-response experiments using concentrations ranging from 1.11 to 30 µM (also lower concentrations were tested for potent compounds). We employed a voltage-ladder protocol to obtain information on SCRA effects on the channel’s voltage dependence of activation (V_50_) and maximal conductance (Gₘₐₓ) (both determined from Boltzmann fits of the instantaneous tail currents; see Methods for details). SCRA efficacy (e.g. maximal reduction in Gₘₐₓ) and potency (SCRA concentration required to produce 50% of the compound’s maximal effect; IC_50_) were determined (see Methods for details). In line with previous reports, JWH-030 inhibited hERG (Fig. 1H-I), seen in our assay as a concentration-dependent decrease in Gₘₐₓ (Fig. 1J, IC_50_ = 23.0 ± 11.8 µM) and a small effect on V_50_ (Fig. 1K). Among the tested compounds, 5F-MDMB-PINACA was one of the more potent inhibitors, reducing Gₘₐ_x_ of hERG at lower concentrations than JWH-030 (Fig. 1J-M, IC_50_ = 9.5 ± 2.5 µM).

A more detailed electrophysiological characterization of the effects of 5F-MDMB-PINACA on hERG is provided in the Supplementary Information (SI Text, SI Fig. S2). Briefly, the compound’s inhibitory effect on hERG was not dependent on temperature or the presence of intracellular fluoride. However, 5F-MDMB-PINACA potency was enhanced in the presence of pluronic acid, likely due to improved availability of the lipophilic 5F-MDMB-PINACA compound. Furthermore, the inhibitory potency was greater in manual patch-clamp (MPC) recordings using adherent cells, whereas recordings from cells in suspension, used both in APC and MPC, yielded similar results. These data highlight the robustness of the 5F-MDMB-PINACA effect under different experimental APC conditions and indicate that our hERG APC assay, if anything, tends to underestimate the potency of SCRAs.

Most SCRAs displayed inhibitory effects on hERG, primarily in the form of decreased Gₘₐₓ (see SI Fig. S3 for concentration-response data and SI Table II for efficacy and potency). For several compounds, the concentration-response curves did not saturate at the highest concentration tested (SI Fig. S3), resulting in suboptimal curve fitting with unreliable efficacy and potency estimates. We therefore compared the effects of the 36 SCRAs at 10 µM, a concentration that provided a consistent and representative estimate of the overall inhibitory effect on hERG. At 10 µM, 24 of the SCRAs significantly reduced Gₘₐₓ (Fig. 1N). In contrast, at 10 µM only 9 of the tested SCRAs induced a significant shift in V_50_ (Fig. 1O). Also, the magnitude of these V_50_ shifts was modest, with all ΔV_50_ confined within a narrow range of ± 7 mV. Hence, for the rest of the study we will focus on the Gₘₐₓ response of hERG to the SCRAs.

### SCRA building block chemistry impacts SCRA inhibition of hERG

We noted that chemical modifications to the building blocks (Fig. 2A) of SCRAs impacted SCRA potency and/or efficacy on hERG (SI Table II). To explore whether broader structural features contribute to these effects we conducted a structure-activity relationship (SAR) analysis both for the efficacy (maximal reduction in Gₘₐₓ) and the potency (IC_50_) of SCRAs on hERG. Only SAR comparisons with at least three pairs of structurally related compounds were included to allow for robust statistical assessment. That the concentration-response curves saturated before full inhibition for several SCRAs (SI Table II, SI Fig. S3) suggested variability in efficacy across SCRAs. However, no consistent SAR patterns could be identified to account for these differences in efficacy (SI Table III-IV). For potency, SCRAs with an indole core were 1.5- to 6.3-fold more potent on hERG than the corresponding SCRAs with an indazole core (Fig. 2B-C, SI Table III). This is exemplified by MDMB-CHMICA vs MDMB-CHMINACA in Figure 2D. Of the different amino acid derived head moieties (Fig. 2E), valinamide (AB) SCRAs were the least potent in all three available comparisons, where tert-leucine methyl esters (MDMB) were 2.6- to 4.1-fold more potent and tert-leucinamides (ADB) were 1.6- to 4.5-fold more potent (Fig. 2F, SI Table IV). There was no difference in potency between MDMB and ADB SCRAs (Fig. 2F, SI Table IV). For tail moieties, no SAR analysis could be performed because of too few structurally related compounds available for comparison or the lack of effect of some compounds (SI Table II).

**Figure 2:**
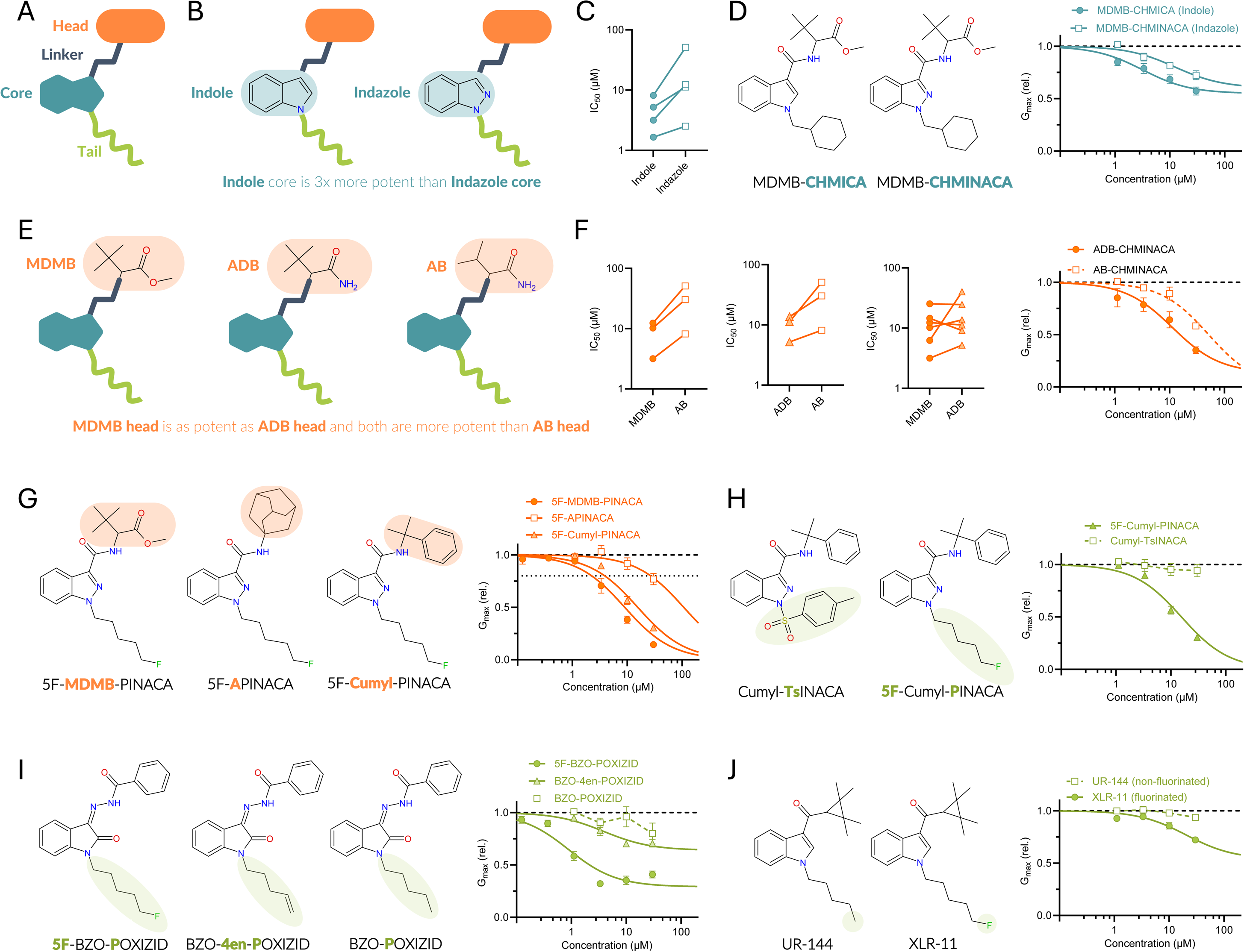
Chemical modifications to the SCRA building blocks impact hERG effects. **(A)** Schematic illustration of the four key building blocks of SCRAs colour coded as in subsequent illustrations. **(B)** Schematic illustration of SCRAs with indole and indazole cores, together with outcome of SAR analysis for hERG potency **(C)**. **(D)** Example of impact of core moiety for hERG inhibition showcased by MDMB-CHMICA and MDMB-CHMINACA. n = 4-5. **(E)** Schematic illustration of SCRAs with indicated heads, together with outcome of SAR analysis for hERG potency **(F)** and example of impact of head moiety showcased by ADB-CHMINACA and AB-CHMINACA for hERG inhibition. n = 4. **(G)** Molecular structures of three SCRAs with 5F-PINACA scaffolds and variable head groups, and concentration-response relationships for indicated SCRAs on hERG inhibition. n = 4-17. **(H)** Molecular structures of two SCRAs with Cumyl-INACA scaffolds and variable tails, and concentration-response relationships for indicated SCRAs on hERG inhibition. n = 4-7. **(I)** Molecular structures of three SCRAs with BZO-OXIZID scaffolds and variable tails, and concentration-response relationships for indicated SCRAs on hERG inhibition. n = 5-12. **(J)** Molecular structures of two SCRAs with variable tails, and concentration-response relationships for indicated SCRAs on hERG inhibition. n = 4-10. All average data are shown as mean ± SEM. Small error bars are covered by symbols. Best fits as in SI Table II.

We also noted several interesting examples of how structural motifs clearly influenced the hERG inhibitory effects of SCRAs, although with too few structurally related compounds to be included in the SAR analysis. For instance, the 5F-PINACA scaffold with either a branched MDMB or aromatic cumyl head group induced concentration-dependent inhibitory effects, whereas the bulky adamantyl in 5F-APINACA impaired the inhibitory effect (Fig. 2G). As another example, a bulky tosyl group at the tail of the Cumyl-INACA scaffold caused Cumyl-TsINACA to be a non-inhibitor (Fig. 2H). Furthermore, addition of a terminal fluorine (5F-pentyl in 5F-BZO-POXIZID) or alkene (pent-4-enyl in BZO-4en-POXIZID) on the pentyl tail of the BZO-POXIZID scaffold produced concentration-dependent inhibitory effects, whereas a simple pentyl tail (in BZO-POXIZID) rendered this SCRA a non-inhibitor (Fig. 2I). As another example of tail chemistry, terminal fluorination of the pentyl tail on the UR-144 scaffold resulted in concentration-dependent inhibitory effects, whereas its non-fluorinated counterpart was a non-inhibitor (Fig. 2J). Similarly, 5F-APINACA showed inhibitory effects, whereas APINACA was a non-inhibitor (SI Table II).

Altogether, the screening of 36 SCRAs on the hERG channel demonstrates that specific structural elements markedly influence hERG inhibition. While some of these features are also reflected in the broader SAR analyses, others are not, underscoring the complexity of SCRA interactions with the hERG channel.

### SCRAs are coordinated in a promiscuous central cavity binding site in hERG

hERG has a known, promiscuous binding site in the central cavity of the pore domain that is used by many inhibitors and blockers ^31,32^, including the histamine receptor antagonist and hERG blocker astemizole ^32–34^ (see also SI Figure S4 for inhibitory effect of astemizole in our hERG assay). This central cavity binding site is just below the selectivity filter and flanked by aromatic residues, like Y652 (schematic illustration in Fig. 3A). To assess whether SCRAs may interact with hERG in this vestibule binding site and predict the mode of interaction, we developed a novel computational pipeline using accelerated weighted histogram simulations, with astemizole as a positive control (SI Fig. S5). The work from the pipeline resulted in the binding poses of astemizole matching the recently solved cryo-EM density ^35^ with astemizole binding perpendicular to the ion conducting pathway in the vestibule binding site, about 5 Å above Y652 (Fig. 3B-C, SI Fig. S5). From this site, astemizole blocks hERG by preventing potassium ion permeation ^35^. We followed the same workflow for four SCRAs, two of which were inhibitors in our hERG assay (5F-MDMB-PINACA and 5F-BZO-POXIZID) and two of which failed to inhibit the channel (BZO-POXIZID and Cumyl-TsINACA). Like astemizole, the simulations with the inhibitors 5F-MDMB-PINACA and 5F-BZO-POXIZID found an energy minimum at 5 Å above Y652 (Fig. 3D-E). Given the time constraint in the AWH simulations, we could not achieve a converged free energy landscape. However, the poses clustered from the energy minima showed the SCRAs in the vestibule in a planar orientation perpendicular to the ion conduction pathway, sandwiched between Y652 and the selectivity filter (Fig. 3F-G for 5F-MDMB-PINACA and 5F-BZO-POXIZID, respectively, and Fig. 3H for a zoomed-out representation of 5F-MDMB-PINACA). This perpendicular conformation is very similar to that of astemizole, thus we propose that this prevents potassium ion permeation. We then took 1000 snapshots from the minima of the cluster to analyse key contacting residues, leading to: i) the identification of Y652 interacting with the SCRA core by pi-pi stacking; ii) S624 and S649 interacting with either the head or the tail of the SCRA, providing hydrogen bonds; and iii) F656 interacting with the alkyl tail by hydrophobic interaction, or defining the size of the head of the SCRAs that can be accommodated in the binding pocket (Fig. 3I). All snapshots from the minima showed the distribution being mostly unimodal, suggesting a very stable binding pose (Fig. 3J-K). Also for the non-inhibitor BZO-POXIZID we found an energy minimum at 5 Å above Y652 (Fig. 3L). However, while we identified a hydrogen bond between the hydroxyl group on S624 and/or S649 and the fluorine of the 5F-pentyl tails of 5F-MDMB-PINACA and 5F-BZO-POXIZID, these are absent for BZO-POXIZID and may explain why it is a non-inhibitor. Hence, although S624 and/or S649 remain in close proximity with the tail in BZO-POXIZID (Fig. 3M), they are unable to form any H-bonding, which is expected to result in a weaker affinity to the channel. Lastly, we sought insight into the structural basis of the non-inhibitor Cumyl-TsINACA. The free energy landscape from the simulations indicates that Cumyl-TsINACA does not show a minimum at 5 Å position (Fig. 3N). This placed both S624 and S649 out of the head binding pocket, misplaced the linker region and disrupted the interaction with the tail (Fig. 3O, with main lost interactions highlighted by *). Together, this suggested how Cumyl-TsINACA fail to block potassium ion permeation.

**Figure 3:**
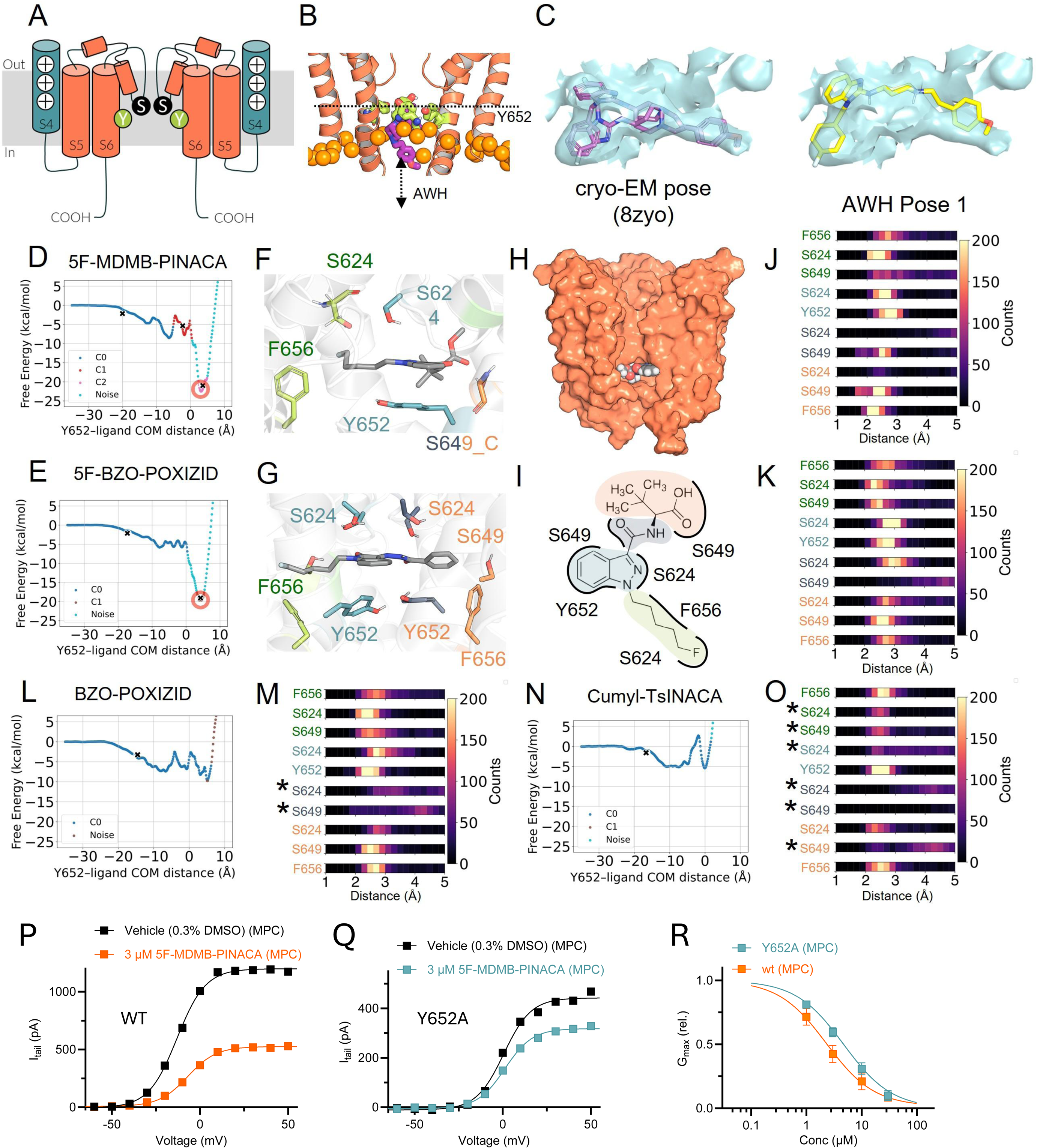
Accelerated weighted histogram simulations provide insight into SCRA binding poses. **(A)** Simplified schematic illustration of hERG (side view) indicating the central ion conducting pathway delineated by the selectivity filter, S624 (denoted S), and Y652 (denoted Y) in the central cavity. Pore domain regions are in orange and the voltage-sensor S4 in blue. **(B)** Structural representation of the path for astemizole (purple) to sample along the z-axis of hERG (orange) with respect to Y652 (green). **(C)** Electron density map obtained from EMD-60574 (cyan) overlayed on the cryo-EM structure of astemizole (purple), comparing to the pose obtained from the minima by AWH (yellow). **(D)** Free energy landscape of the hERG inhibitor 5F-MDMB-PINACA sampling along the z-axis with respect to Y652, starting from the docking pose (see methods for details). Red circle and a cross mark the minima of the landscape. Different colours indicate different cluster extracted from DBSCAN with the epsilon = 0.2 with at least 10 frames in the cluster. **(E)** Same as in D but for 5F-BZO-POXIZID. **(F-G)** Snapshots of 5F-MDMB-PINACA (F) and 5F-BZO-POXIZID (G) from the cluster centre of the free energy minima. The C-atoms of the SCRA are coloured grey. Residues interacting with the SCRAs are coloured according to which building block they interact with, with the head group in orange, the linker in black, the core in blue and the tail in green (same colour coding as in Figure 2). **(H)** A zoom out picture of SCRA binding site on the pore domain of hERG (shown as orange surface). 5F-MDMB-PINACA is shown as a black sphere with CPK colouring for the elements. **(I)** Schematic representation of SCRAs in the binding pocket (here illustrated for 5F-MDMB-PINACA), forming interactions with residues on hERG subunits. (**J-K**) Distribution of minimum distance analysis from 1000 structures at the minima performed between each residue in four hERG subunits and the four building blocks of the SCRAs for 5F-MDMB-PINACA (J) and 5F-BZO-POXIZID (K). Residue labels on the left side are colour-coded according to the building block they contact (same colours as in panels F and G). The number of contact each interaction is shown on the colour bar to the right side. The bin size of the histogram is 20. **(L-M)** Same as in panels D and J but for the non-inhibitor BZO-POXIZID. **(N-O)** Same as in panels D and J but for the non-inhibitor Cumyl-TsINACA. The asterisks mark residues with no contact. **(P-Q)** Representative effect of 3 µM 5F-MDMB-PINACA on hERG wild type (WT) (P, stable cell line) or hERG Y652A (Q, transient cell line) collected using MPC. **(R)** Concentration-response relationship for the G_max_ (rel) effect of 5F-MDMB-PINACA on hERG WT (orange) and Y652A (teal), collected using MPC. Best fits for G_max_ (rel) yielded IC_50_ values of 2.3 µM (n = 5–13) for WT and 4.7 µM (n = 8–9) for Y652A. The IC_50_ values were significantly different (p < 0.05) using Extra sum-of-squares F-test. For both WT and Y652A, efficacy approached full inhibition.

By mutating Y652 to alanine (Y652A), the proposed pi-stacking interaction between the compound’s core and the residue is disrupted. We compared the 5F-MDMB-PINACA effect on hERG wild type and the Y652A mutant. 5F-MDMB-PINACA was less potent on the Y652 mutant (Fig. 3P-R), as if Y652 is one of the residues involved in 5F-MDMB-PINACA binding or effects. However, despite the reduced potency, the 5F-MDMB-PINACA still exerts an inhibitory effect, likely due to preserved or compensatory interactions with other residues. The reduced potency of 5F-MDMB-PINACA induced by Y652A is in the range of that previously reported for the hERG inhibitor m-hydroxymexiletine ^36^, but smaller than that of hERG inhibitors like amiodarone and astemizole ^37^.

Altogether, we conclude that for SCRAs to assume a mode of hERG interaction in the central cavity preventing potassium ion permeation, we postulate two rules: (i) The head group and the tail must be small enough to fit in the binding pocket dictated by F656. A larger head group (e.g. adamantyl) or tail group (e.g. tosyl) would prevent the SCRA from binding properly, resulting in a shift in the position of the minima in the free energy landscape. (ii) Either the head group (MDMB or Cumyl) or the fluorinated tail must be able to form hydrogen bonds with S624 or S649. SCRAs with no tail, such as MDMB-5’Br-INACA or ADB-5’Br-INACA will result in a weaker interaction with the binding pocket. Given the limitation on the convergence of the free energy in the simulations, it is challenging to determine how the small differences between the indole and indazole cores impact the pi-stacking interaction with Y652. However, indazole has been shown to be less electron-rich than indole due to the additional p-orbital on the N atom ^38^. This may weaken the pi-stacking interactions with Y652.

### Comparing SARs for SCRA effects on hERG and the CB_1_ receptor reveal similarities and differences

To explore the relationship between the potential cardiotoxic effects of SCRAs via hERG and their psychoactive effects mediated by the CB_1_ receptor, we assessed the activity of the 36 SCRAs on CB_1_ stably expressed in CHO-K1 cells using an AequoScreen^®^ intracellular calcium release assay (see Methods for details; SI Fig. S6A for exemplary example of JWH-018). The compounds showed a wide range of potencies on CB_1_, with EC_50_ values spanning from sub-nanomolar (e.g., MDMB-FUBINACA) to low micromolar levels (e.g., BZO-HEXOXIZID) (Fig. 4A).

**Figure 4.**
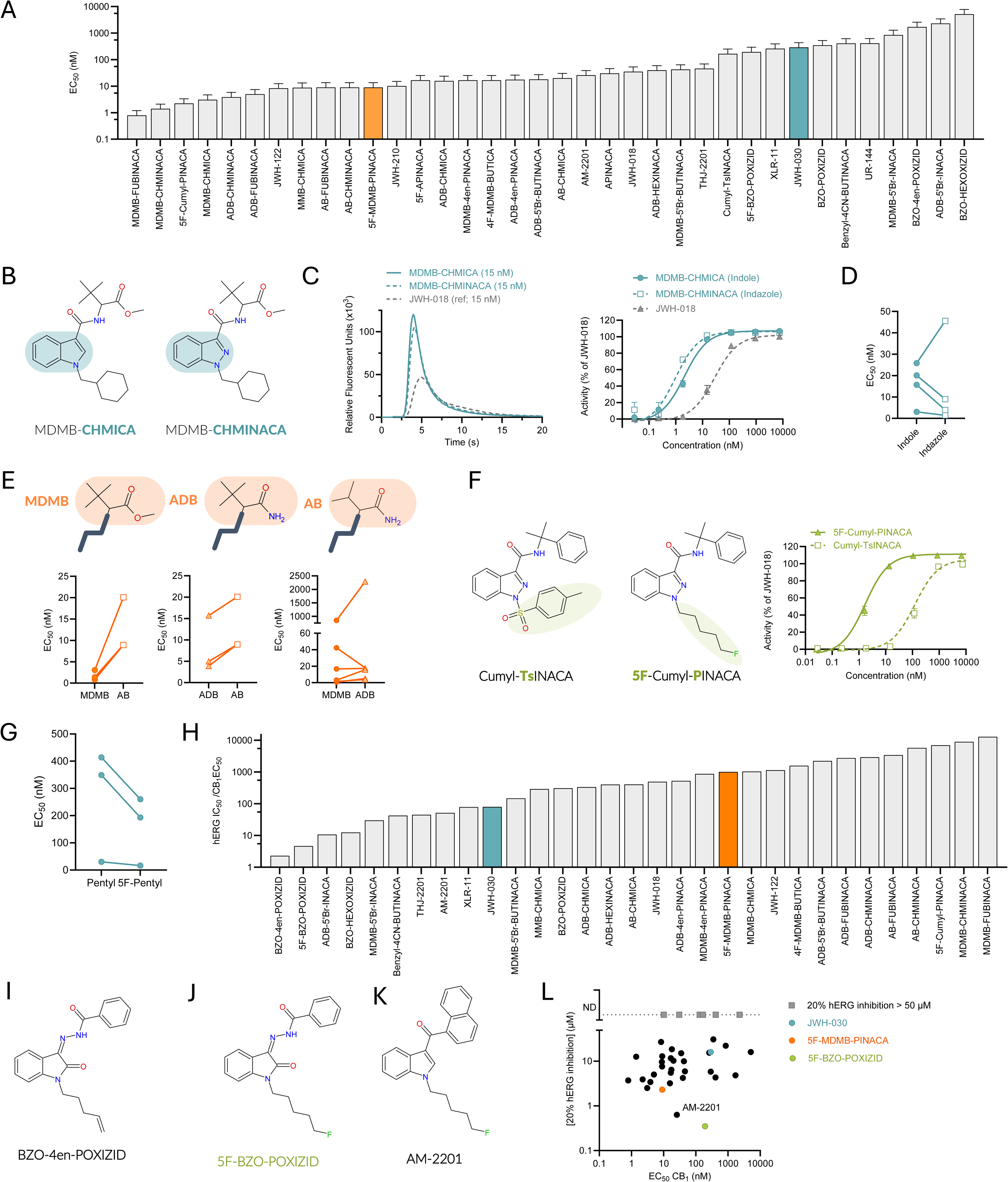
Comparison of SCRA features important for CB_1_ and hERG effects reveal similarities and differences dictating SCRA potency. **(A)** Potency of the indicated SCRAs on CB_1_ (see methods for details). **(B-C)** Example of impact of core moiety for CB_1_ activation showcased by MDMB-CHMICA and MDMB-CHMINACA (structures in B and CB_1_ activating effects in C). **(D)** Outcome of SAR analysis of SCRAs with indole and indazole cores for CB_1_ potency. **(E)** Schematic illustration of SCRAs with indicated heads, together with outcome of SAR analysis for CB_1_ potency. **(F)** Molecular structures of two SCRAs with Cumyl-INACA scaffolds and variable tails, and concentration-response relationships for indicated SCRAs on CB_1_ activity. **(G)** Outcome of SAR analysis of SCRAs with non-fluorinated compared with fluorinated pentyl tails for CB_1_ potency. **(H)** Ranking of the cardiotoxic harm potential of the SCRAs based on the ratio of hERG IC_50_ to CB_1_ EC_50_, where a low ratio indicates higher cardiotoxic harm. SCRAs where the hERG IC_50_ value was unable to be calculated are not included in this figure. **(I-K)** Molecular structure of indicated SCRAs. **(L)** Plot of SCRA concentration causing 20% inhibition of hERG versus CB_1_ EC_50_. All average data are shown as mean ± SEM. Small error bars are covered by symbols. Data in panels H and L are shown as mean only due to the nature of this data. For CB_1_ data is from a minimum of three independent experiments (n ≥ 3), run in triplicate (except for JWH-030, for which n = 2). n for hERG and best fits as in SI Table II.

In contrast to the results on hERG, MDMB-CHMICA and MDMB-CHMINACA showed similar potency on CB_1_ (Fig. 4B-C, SI Table V). Moreover, no consistent pattern in CB_1_ potency was observed between compounds with indole or indazole cores (Fig. 4D). For the head moiety, a similar overall SAR was seen for CB_1_ receptor activity as for hERG, where tert-leucine methyl esters (MDMB) and tert-leucinamides (ADB) were consistently more potent than the corresponding valinamide (AB) head moiety (Fig. 4E, SI Table VI). Moreover, an adamantyl group (5F-APINACA) instead of a MDMB or cumyl group reduced the hERG inhibitory effect and also reduced the CB_1_ potency (SI Fig. S6B).

Similar to the results on hERG, replacing the fluorinated pentyl tail of 5F-Cumyl-PINACA with the tosyl tail of Cumyl-TsINACA reduced the potency at the CB_1_ receptor, shifting the EC_50_ from 1.8 nM to 123 nM (Fig. 4F). For hERG, the importance of fluorination was further supported by the increased potency of 5F-BZO-POXIZID compared to BZO-POXIZID, and similarly for UR-144 versus XLR-11. At the CB_1_ receptor, a similar importance of tail fluorination was seen, as fluorinated SCRAs tended to show higher potency compared to their non-fluorinated counterparts (Fig. 4A, G, SI Table VII). Taken together, certain structural features, particularly in the head and tail moieties, show similarities in how they contribute to effects in hERG and CB_1_, whereas the core scaffold appears to contribute differently to activity on the two targets. Therefore, the potency of SCRAs on hERG and CB_1_ show little apparent association (SI Fig. S6C), reflecting distinct SARs.

### Comparing hERG and CB_1_ potency ratios and effects for risk stratification

Assessing the relative risk profile of SCRAs remains complex due to the multifaceted nature of their pharmacology. Based on our data, one approach to evaluating the potential for cardiotoxicity is to consider the relative potency at the CB_1_ receptor compared to the hERG channel, using the ratio of hERG IC_50_ to CB_1_ EC_50_. Since CB_1_ potency reflects psychoactive strength, SCRAs with high hERG potency but low CB_1_ potency (i.e., a low ratio) may pose a greater cardiotoxic risk, as higher doses are likely required to achieve the desired psychoactive effects. The ratio of hERG IC_50_ to CB_1_ EC_50_ varied from 2.3 to over 12,000 for the 36 SCRAs we studied (Fig. 4H). All SCRAs had a ratio >1, indicating that a greater concentration of the SCRA is required to inhibit hERG than to achieve psychoactive effects (although, comparing absolute concentrations between different assays is a challenge). By this standard, BZO-4en-POXIZID (Fig. 4I) had the highest risk of cardiotoxicity, i.e., the lowest ratio. 5F-BZO-POXIZID (Fig. 4J) is also connected to a high risk of cardiotoxicity, with only a five-fold higher potency of CB_1_ compared to hERG (Fig. 4H). In general, the OXIZID SCRAs exhibited a high cardiotoxic risk, whereas the CHMINACA and FUBINACA SCRAs had a comparatively low cardiotoxic risk.

The pro-arrhythmic potential of other classes of compounds inhibiting hERG has been extensively studied in the past, for which compounds inducing hERG inhibition by 20% have been indicated as ‘risky’ compounds likely to induce detectable changes to action potential duration and QT interval ^39,40^. Given that some SCRAs showed low efficacy as hERG inhibitors (SI Table II), another way of evaluating the relative potential for the greatest cardiotoxicity of SCRAs is based on the potency at the CB_1_ receptor versus the SCRA concentration inducing 20% reduction in hERG conductance. In this regard, SCRAs like AM-2201 (Fig. 4K), with low CB_1_ receptor potency combined with relatively low concentration required to cause 20% hERG inhibition, are particularly concerning (Fig. 4L). Based on our data, it is notable that several SCRAs show similar risk profiles as two of the SCRAs with previously established clear cardiotoxic effects and/or hERG inhibition and action potential prolonging capability (5F-MDMB-PINACA and JWH-030) (Fig. 4L).

### Several SCRAs show dual inhibitory effects by targeting both hERG and K_V_7.1/KCNE1

Both hERG and K_V_7.1/KCNE1 contribute to cardiomyocyte repolarization (Fig. 1G). Inhibition of K_V_7.1/KCNE1 alone can be proarrhythmic ^41^. Moreover, under conditions with impaired hERG function, upregulation of K_V_7.1/KCNE1 has been proposed to partially compensate for the loss of hERG ^42,43^. Hence, drugs that inhibit both hERG and K_V_7.1/KCNE1 may have compounded proarrhythmic effects ^44^, which is another factor to consider when evaluating potential cardiotoxic effects of SCRAs. To assess if SCRAs also affect K_V_7.1/KCNE1, we used mammalian CHO cells stably expressing human K_V_7.1/KCNE1. Several SCRAs displayed inhibitory effects on K_V_7.1/KCNE1, seen as reduced G_max_ and/or shifted V_50_ (SI Fig. S7-8, SI Table II). A notable example was MDMB-5’Br-BUTINACA (Fig. 5A), which reduced Gₘₐₓ and shifted V_50_ towards more positive voltages (Fig. 5B-D). Both inhibitory effects on K_V_7.1/KCNE1 were also seen for other SCRAs, including 5F-Cumyl-PINACA, MDMB-CHMINACA, MDMB-4en-PINACA and THJ-2201 (SI Fig. S7-8). At 10 µM, 7 of the SCRAs caused a significant positive shift in V_50_ of activation (Fig. 5E), with the largest shift induced by MDMB-CHMINACA (ΔV_50_ = +10.3 ± 1.6 mV). At 10 µM, 9 of the SCRAs significantly reduced Gₘₐₓ (Fig. 5F). In contrast to hERG, 5F-MDMB-PINACA had a small effect on K_V_7.1/KCNE1, while JWH-030 induced the largest reduction (Fig. 5E-I, SI Table II). Hence, several SCRAs inhibited K_V_7.1/KCNE1, however, they were fewer and induced less prominent reduction in Gₘₐₓ compared to what was observed for hERG.

**Figure 5.**
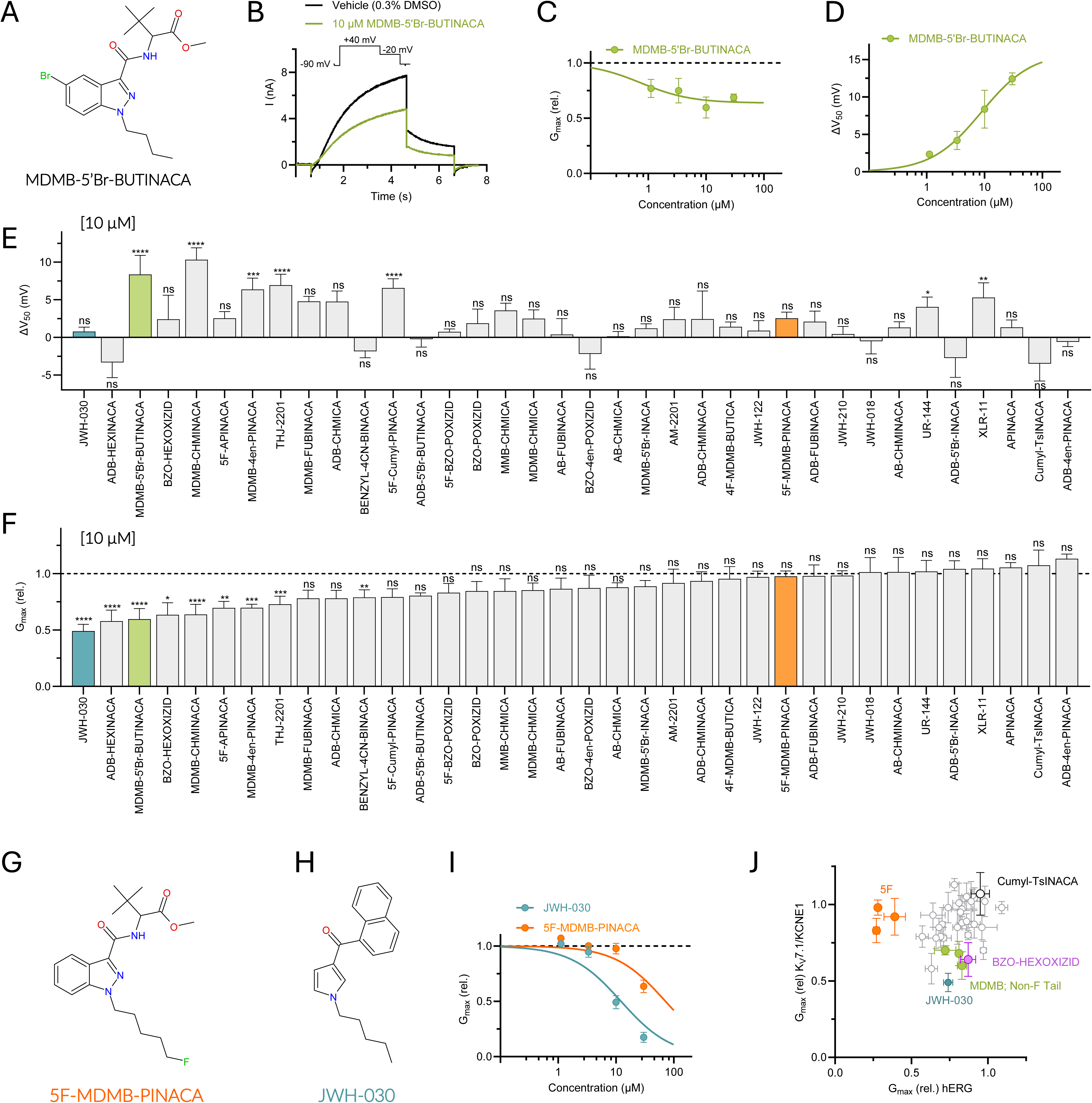
Some tested SCRAs inhibit also K_V_7.1/KCNE1. **(A)** Chemical structure of MDMB-5’Br-BUTINACA. **(B)** Representative effect of MDMB-5’Br-BUTINACA on K_V_7.1/KCNE1 using the pulse protocol shown as insert. **(C-D)** Concentration-response relationship for the rel. G_max_ (C) or V_50_ (D) effect of MDMB-5’Br-BUTINACA on K_V_7.1/KCNE1. n = 4-5. **(E-F)** Average effect of 10 µM of the indicated SCRAs on V_50_ (E) and G_max_ (F) of K_V_7.1/KCNE1. n = 4-9. Statistics denote adjusted p-values for comparisons to vehicle: p < 0.05 (*), p < 0.01 (**), p < 0.001 (***), p < 0.0001 (****). **(G-I)** Chemical structure of 5F-MDMB-PINACA (G) and JWH-030 (H) with concentration-response relationship for their rel. G_max_ effects on K_V_7.1/KCNE1. (I). n = 6. **(J)** Comparison of G_MAX_ response of K_V_7.1/KCNE1 (y-axis) and hERG (x-axis) induced by SCRAs at 10 µM. Specific SCRAs are highlighted as indicated in the figure. Three SCRAs with 5F-pentyl tails (5F-MDMB-PINACA, 5F-BZO-POXIZID, AM-2201) are indicated in orange. Three SCRAs with MDMB heads and non-fluorinated tails (MDMB-5’Br-BUTINACA, MDMB-4en-PINACA, MDMB-CHMINACA) are indicated in green. Note that SCRAs marked in grey were either non-inhibitors of hERG or had an anticipated IC_50_ exceeding 50 µM. All average data are shown as mean ± SEM. Small error bars are covered by symbols. Best fits and n as in SI Table II.

Given the limited number of SCRAs with effects on K_V_7.1/KCNE1 no SAR analysis could be performed. However, we note several interesting clusters of compounds, exemplified for 10 µM in Figure 5J. At 10 µM, some SCRAs had no effect on either of the two channels (e.g. Cumyl-TsINACA in black), whereas some had effects only on hERG (e.g. 5F-MDMB-PINACA, 5F-BZO-POXIZID, and AM-2201 in orange) or only on K_V_7.1/KCNE1 (e.g. BZO-HEXOXIZID in purple). Yet other SCRAs had inhibitory effects on both hERG and K_V_7.1/KCNE1 (e.g. JWH-030 in blue and MDMB-5’Br-BUTINACA, MDMB-4en-PINACA, and MDMB-CHMINACA in green), showing compounded inhibitory effects on both these repolarizing K_V_ channels.

## Discussion

We demonstrate that the majority of SCRAs included in our study inhibit hERG, primarily by reducing the channel conductance. However, the magnitude of the effects and concentrations at which they occur varies among SCRAs. Generally, we noted that SCRAs with an indole core are more potent hERG inhibitors than their indazole core counterparts. Moreover, we observed important roles for both the head and tail groups. For instance, we found that several SCRAs with fluorinated alkyl tails (mostly 5F-pentyl) are prominent hERG inhibitors. This should be a cause for concern, as 5F-pentyl is the second most common tail moiety among SCRAs ^45^. Moreover, bulky and inflexible groups often impaired hERG effects, showcased by Cumyl-TsINACA (with an inflexible, bulky tosyl tail) and 5F-APINACA (with a bulky and inflexible adamantyl head group). Our data suggest that SCRAs use the canonical hERG central cavity binding pocket, adopting a planar orientation perpendicular to the ion conducting pathway. Interestingly, we found only partial correlation between how SCRA structural features impact the potency to inhibit hERG compared to activating the CB_1_ receptor. Moreover, inhibitory SCRA effects on hERG and K_V_7.1/KCNE1 showed no apparent correlation, as fewer SCRAs had inhibitory effects on K_V_7.1/KCNE1 compared to hERG, with some compounds affecting only one of the channels. These findings indicate that different chemical properties are important for SCRA binding or effects on these different targets and demonstrate the need to experimentally test the effect of SCRAs on multiple targets.

The SCRA orientation seen in our simulations is very similar to that of cryo-EM structures of the hERG pore blockers astemizole, E-4031 and pimozide ^35^, and the K_Ca_3.1 pore blocker 1,4-dihydropyridine ^46^, which all are shown to lie flat below the selectivity filter, form interaction with aromatic side chains, and are proposed to prevent ion permeation. Hence, although alternative binding poses and mechanisms underlying hERG inhibition by SCRAs are possible, we find it likely that SCRAs like 5F-MDMB-PINACA and 5F-BZO-POXIZID act as hERG pore blockers by obstructing and preventing currents through hERG. The susceptibility of hERG to SCRAs can be explained by the coordination geometry in the binding pocket, where Y652 provides pi-stacking, and F656 dictates the size and the shape of the head or the tail. In addition, S624 and S649 provide hydrogen bonding partners to the SCRAs, tuning the affinity to the different functional groups. In contrast, our SCRA orientation is different from that recently proposed for 5F-APINACA, which uses a side pocket lateral to the central cavity ^27^. One possibility is that a more lateral binding site is preferred by SCRAs that do not coordinate well in the central cavity, which would agree with our observation that a bulky adamantyl group (as in 5F-APINACA) is expected to impair SCRA interaction in the central cavity. That SCRAs can interact with lateral sites may also contribute to the lack of complete hERG inhibition at saturating concentrations for some SCRAs.

In modern structural biology, cryo-EM is a common tool for determining the drug-binding pocket and its interaction geometry. Information from the cryo-EM density map can be an excellent starting point for identifying the binding pose of related drug molecules, or even for understanding how they gain access to the binding site. To enrich detailed cryo-EM determined drug binding sites, computational approaches may reveal the dynamics of drug binding and drug pose adaptation. Well-tempered metadynamics recently identified a binding site of the Na_V_1.5 channel blocker flecainide in the pore just below the selectivity filter. This was very close to the site of flecainide in the cryo-EM structure, and also revealed how different compounds prefer either the central cavity or lateral fenestrations of Na_V_1.5 ^47^. In our study, we used accelerated weighted histogram simulations, which is similar to metadynamics but is native to GROMACS ^48^, to determine the binding of astemizole to the hERG channel with a binding mode matching the electron density map. Moreover, we extended our computational approach to four SCRAs, showing that the shape of the free energy landscape can be used to determine the efficiency of the blocker. Hence, in addition to providing mechanistic insights into SCRA effects, our study highlights accelerated weighted histogram simulations, just like metadynamics, as a valuable tool in providing insights into the binding dynamics and versatility of ion channel inhibitors targeting the ion permeation pathway.

How do our findings on hERG inhibition by SCRAs compare with the current literature? Yun *et al.* reported JWH-030 to be an inhibitor of hERG capable of inducing QT prolongation in anesthetized rats ^26^; however, their paper reports an IC_50_ of 88.4 µM (4-fold greater than our IC_50_ of 21.3 ± 11.9 µM). Another study employed machine learning tools to predict hERG interaction of seven SCRAs ^28^. Our electrophysiology data agree with their predictions for ADB-4en-PINACA and ADB-5’Br-INACA, showing no or limited effects for these SCRAs, and for 5F-BZO-POXIZID, showing prominent inhibitory effects in our hERG assay. In a more recent study, Cheng *et al.* reported inhibitory hERG effects of 5F-APINACA with an IC_50_ of 2.2 µM ^27^, which is considerably lower than our IC_50_ (>100 µM, SI Table II). Hence, our study agrees with these reports in observing inhibitory SCRA-mediated effects on hERG, whereas experimental differences may contribute to the varying observed concentrations required to achieve effects. To our knowledge, no additional SCRA has been functionally tested on hERG, but reports on the observed side effects of the use of some SCRAs are consistent with our findings. MDMB-CHMICA has been reported in two cases of sudden cardiac death ^49,50^, and multiple analytically confirmed intoxications have reported tachycardia and ECG changes ^51,52^. Our data show that MDMB-CHMICA was one of the SCRAs requiring the lowest concentration to inhibit hERG by 20%. Moreover, 5F-MDMB-PINACA is reportedly one of the most dangerous SCRAs, linked to several fatalities ^18–20,53,54^ and non-fatal intoxication with symptoms suggestive of cardiac arrhythmia (palpitations, heart pain, syncope and anxiety) ^21^. We found this SCRA to be one of the most prominent inhibitors of hERG, with high efficacy, relatively high potency, and a low concentration needed to inhibit hERG by 20%.

Besides isolated effects on hERG, other aspects can be considered when aiming to understand links between SCRAs and adverse cardiac events. One such aspect is the relative effect on hERG compared to the CB_1_ receptor, where SCRAs requiring high concentrations to activate CB_1_ are of higher risk of inducing adverse effects via other targets. In our experiments, we note that OXIZID and tail-less SCRAs show the most unfavourable CB_1_ to hERG concentration relationship. It is worth noting, though, that halogenated tails might limit the penetration over the blood brain barrier and hence access to CB_1 55_, which would contribute to a more dangerous risk profile as higher peripheral SCRA concentrations would be required to achieve central CB_1_ effects. Additionally, several SCRAs also inhibited K_V_7.1/KCNE1 in our experiments, which is anticipated to induce dual detrimental effects on cardiac repolarization. Hence, based on our findings exploring SCRA effects on hERG, K_V_7.1/KCNE1 and CB_1_, we reason that SCRA-induced inhibition of hERG alone, or both hERG and K_V_7.1/KCNE1, can contribute to cardiotoxic effects, should the local SCRA concentration in the heart reach sufficiently high concentrations. Other factors, including polydrug intoxication ^20,56^, congenital predisposition ^23,57^ or electrolyte imbalance ^51,52,58,59^, may contribute to cardiac adverse effects following SCRA use.

Study limitations include difficulties in comparing SCRA potency across different targets, due to the use of multiple assays and SCRA delivery methods, combined with SCRAs’ high lipophilicity ^55^. This also contributes to challenges in relating our findings to relevant *in vivo* concentrations, further complicated by a lack of accurate dosing information for SCRAs (as it is only available in most cases based on self-reports by those using SCRAs). The concentration of some SCRAs have been detected in post-mortem cases. For instance, the concentration of 5F-MDMB-PINACA in post-mortem cases are reported to be about 2 ng/mL (approx. 5.3 nM) ^18–20,53,54^ in blood or serum samples, and 1.82 ng/mL (approx. 4.8 nM) in the heart muscle ^60^. However, it is not possible to interpret how well those concentrations reflect those reached in people who use SCRAs, or the original dose used. Another study limitation is that the SCRA library would need to be expanded beyond our 36 SCRAs for additional SAR analysis.

In conclusion, our extensive functional screening of SCRAs found multiple compounds with inhibitory actions on one or both repolarizing cardiac K_V_ channels, with several of the compounds linked to cardiotoxic adverse events in literature. Our study lends support to a central cavity hERG binding site for several SCRAs, highlights chemical motifs tuning the SCRA effect on hERG, while also providing insights into similarities and differences in SCRA features impacting CB_1_ and K_V_7.1/KCNE1 effects. While this screen represents 13% of the 277 SCRAs that are monitored by the EUDA (formerly the EMCDDA) as of 2025 ^61^, our study not only provides mechanistic insights into adverse effects of SCRAs but also motivates the evaluation of ion channel effects and cardiac effects of more SCRAs, adding another dimension to the safety and risk assessment of new drugs detected on the recreational drug market.

## Methods

### Chemicals

4F-MDMB-BUTICA, 5F-APINACA, ADB-4en-PINACA, ADB-5’Br-BUTINACA, ADB-CHMICA, ADB-FUBINACA, ADB-HEXINACA, Benzyl-4CN-BUTINACA, Cumyl-TsINACA, JWH-210, MDMB-CHMICA, MDMB-CHMINACA, MDMB-FUBINACA, and THJ-2201 reference standards (purity ≥ 98%) were purchased from Cayman Chemicals (Ann Arbor, USA). 5F-BZO-POXIZID, 5F-Cumyl-PINACA, 5F-MDMB-PINACA, AB-CHMICA, AB-CHMINACA, AB-FUBINACA, ADB-5’Br-INACA, ADB-CHMINACA, AM-2201, APINACA, BZO-POXIZID, BZO-4en-POXIZID, BZO-HEXOXIZID, JWH-018, JWH-030, JWH-122, MDMB-4en-PINACA, MDMB-5’Br-INACA, MDMB-5’Br-BUTINACA, MMB-CHMICA, UR-144, and XLR-11 reference standards (purity ≥ 98%) were purchased from Chiron AS (Trondheim, Norway). Astemizole was purchased from Sigma-Aldrich (Stockholm, Sweden). All compound solutions were in DMSO at a concentration of 10 mM and stored at -20 °C. Compounds were diluted in extracellular recording solution on the day of recordings. For the automated patch-clamp experiments, the compounds made no contact with plastics following dilution in extracellular solution. The final concentration of DMSO was equal in all recording solutions to control for vehicle effects. For experiments involving pluronic acid (Pluronic F-68; P5556; POLOXAMER 188 SOLUTION; Merck Life Science AB, Sweden), compound stock solutions were diluted in 2 mM pluronic acid (10% solution), warmed to approximately 60°C, and sonicated for 30 seconds. The final concentration of pluronic acid in the assay was 0.05%. DMEM/Ham’s F12, Ham’s F12, 0.25% trypsin-EDTA with phenol red, and fetal bovine serum (FBS) were purchased from Thermo Fisher (Gothenburg, Sweden). HEPES buffer, L-glutamine, protease-free bovine serum albumin (BSA), digitonin, adenosine-5’-triphosphate disodium salt hydrate (ATP), methanol, and DMSO were procured from Sigma-Aldrich (Stockholm, Sweden). The coelenterazine substrate was from Nanolight Technology (Pinetop, AZ, United States).

### Cell lines

Stably expressing cell lines “CHO-hERG DUO” (here referred to as hERG; gene: KCNH2) and “CHO-K_V_LQT1/mink” (here referred to as K_V_7.1/KCNE1; gene: KCNQ1/KCNE1), (B’SYS GmBH, Witterswil, Switzerland) were used to study whole-cell K^+^ currents on a QPatch II 48 APC system (Sophion Bioscience A/S, Ballerup, Denmark). Cell culture and passaging was performed in accordance with standard operating procedures. Cells were harvested in serum-free medium with Detachin^TM^ (Genlantis, CA, USA) immediately prior to experiments. For MPC on transient cell lines, CHO-K1 CCL-61 cells (ATCC, Manassas, VA, USA) were cultured in Ham’s F-12 medium supplemented with 10% fetal bovine serum (FBS) and 100 U/mL penicillin–100 µg/mL streptomycin (Gibco, Thermo Fisher Scientific, Waltham, MA, USA). Transfection was carried out using JetOPTIMUS® reagent (Polyplus, Strasbourg, France) according to the manufacturer’s protocol. Briefly, 80,000 cells were seeded per well in a 12-well plate and transfected with 300 ng of pcDNA3-GFP plasmid, 700 ng of pcDNA3-hERG plasmid with the Y652A mutation, and 1 µg of JetOPTIMUS. After 8 hours, cells were detached and reseeded at a density of 30,000 cells per well onto several poly-D-lysine-coated glass coverslips placed in a 12-well plate. For the CB_1_ receptor activity assay, the AequoScreen® recombinant Chinese hamster ovary (CHO) K1 cell line, stably expressing the human CB_1_ receptor (ES-110-A), was used (Revvity, Sollentuna, Sweden).

### Automated patch-clamp experiments

Automated patch-clamp recordings were performed at 22°C, if not else indicated. Freshly harvested cell suspensions (2-3 million cells/mL) were prepared for experiments by the QPatch II 48 automated cell preparation unit. Cells were centrifuged for 150 s at 150 x g and washed twice with the extracellular recording solution. Cells were handled by the integrated pipetting robot and added to the 48 experiment sites on the planar patch-clamp consumable QPlate. Each experiment site contained an individual pair of recording electrodes and a silicon/glass biochip embedded in the flow channels. Single-cell patch recordings and population patch recordings (10 cells per well, n = 1 is defined as one population patch) were performed for hERG and for K_V_7.1/KCNE1, respectively. Cells were applied, positioned and sealed to the patch holes by a mild suction protocol; whole-cell access was then achieved by a stronger suction pulse.

The extracellular solution contained (in mM): 145 NaCl, 10 Glucose, 4 KCl, 2 CaCl_2_, 1 MgCl_2_, and 10 HEPES. Osmolarity was adjusted to 305-308 mOsm with sucrose and pH was adjusted to 7.4 with NaOH. The intracellular solution contained (in mM): 100 KCl, 4.3 CaCl_2_, 1.4 MgCl_2_, 25/10 KOH/EGTA, 24 KF, 10 HEPES, and 4 Mg-ATP (hERG) or 4 Na-ATP (K_V_7.1/KCNE1). Osmolarity was adjusted to 305-308 mOsm with sucrose, and pH was adjusted to 7.2 with KOH.

Square voltage pulses in a voltage ladder protocol were used to study current/voltage (I(V)) relationships. Each voltage protocol was first run three times in the presence of vehicle (extracellular solution supplemented with 0.3% DMSO), the 3^rd^ serving as the baseline. Voltage protocols were repeated in the presence of 4 cumulatively rising concentrations of test compound, followed by application of vehicle for recovery. To control for assay quality, one set of vehicle-only experiments which received no test compound was included in each run. Each solution was added to the QPlate twice (5 s between applications: 10 µL per application). For the hERG voltage ladder protocol, solutions were present for 60 s before recording started. hERG cells were kept at a holding voltage of −80 mV, with brief hyperpolarizing steps to −90 mV for 50 ms (to calculate ohmic leakage) before returning to −80 mV for 200 ms. Square voltage pulses were sequentially applied in 10 mV increments (range −60 mV to +60 mV) for 5 s before stepping to −50 mV for 2 s to record tail currents. Time between start of sweeps was 10 s. The voltage pulsing protocol was executed every 150 s. In experiments using the hERG voltage pulsing protocol, the extracellular solution remained present for 7 s before recording started. hERG cells were kept at a holding voltage of −80 mV with brief hyperpolarizing steps to −90 mV for 50 ms (to calculate ohmic leakage) before returning to −80 mV. Stepping occurred in two stages, first to −50 mV for 50 ms before stepping up +40 mV for 5 s. Cells were then held −50 mV for 2 s to record tail currents and returned to −80 mV. Time between start of sweeps was 10 s, and sweeps were pulsed 20 times per protocol. The voltage pulsing protocol was executed every 201 s.

In experiments with fluoride-free intracellular solution (SI Fig S2F-G), the intracellular solution contained (in mM) 120 KCl, 5.374 CaCl_2_, 1.75 MgCl_2_, 31.25/10 KOH/EGTA, 10 HEPES, and 4 Na-ATP. Osmolarity was adjusted to 285-295 mOsm with sucrose, and pH was adjusted to 7.2 with KOH. Extracellular solution (5 µl) was added to the recording plate, and the solution remained present for 9 s before recording started. hERG cells were kept at a holding voltage of −90 mV with brief hyperpolarizing step from −100 mV for 100 ms to −110 mV for 100 ms (to calculate ohmic leakage) before a brief step to −50 mV for 100 ms (to enable automated leak subtraction) followed by an activating step to +20 mV for 4 s. Cells were then held at −50 mV for 4s to record tail currents and returned to −100 mV for 1 s. Time between start of sweeps was 15 s, and sweeps were pulsed 10 times. Following the pulsing protocol, extracellular solution (5 µl) was added to the recording plate, and the solution remained present for 5 s before voltage-ladder protocol started with a brief hyperpolarizing step from −100 mV for 100 ms to −110 mV for 100 ms (to calculate ohmic leakage) before a brief step to −50 mV for 100 ms (to enable automated leak subtraction). Square voltage pulses were sequentially applied in 10 mV increments (range −100 mV to +70 mV) for 4 s before stepping to −50 mV for 4 s to record tail currents, followed by a step to −100 mV for 1 s. Time between start of sweeps was 9.32 s, and the voltage ladder protocol was executed every 170 s. Voltage ladder protocols were repeated in the presence of 2 cumulatively rising concentrations of test compound: 2 µl was added with 5 repetitions (2 s between each application) followed by application of extracellular solution (2 µl was added with 5 repetitions (2 s between each application)) for recordings of recovery using the precedent voltage pulsing protocol. The temperature was set to 27°C.

For the K_V_7.1/KCNE1 voltage ladder protocol, the solutions were present for 60 s before recording started. K_V_7.1/KCNE1 cells were kept at a holding voltage of −90 mV, with brief hyperpolarizing steps to −100 mV for 540 ms (to calculate ohmic leakage). Square voltage pulses were sequentially applied in 10 mV increments (range −100 mV to +40 mV) for 4 s before stepping to −20 mV for 2 s to record tail currents. Time between compound application and first voltage protocol was 30 s. Time between sweeps was 15 s. The voltage ladder protocol was executed every 220 s.

### Transient transfections and manual patch-clamp electrophysiology

Manual patch-clamp recordings were performed 24–48 hours post-transfection. Coverslips with CHO-K1 cells were placed in a recording chamber and continuously perfused at room temperature (22–24⁰C) with extracellular solution by a gravity-fed perfusion system.

Compounds were applied by a pressurized, automated OctaFlow perfusion system (ALA Scientific Instruments). The intracellular and extracellular solutions used MPC were identical to those used for APC electrophysiology (see above). Patch pipettes were pulled from borosilicate glass and had resistances of 3–6 MΩ when filled with intracellular solution.

Square voltage pulses were sequentially applied in 10 mV increments (range −60 mV to +50 mV from a holding potential of −80 mV) for 5 s before stepping to −50 mV for 2 s to record tail currents. Each voltage protocol was first run three times in extracellular solution, the 3^rd^ serving as the baseline. Voltage protocols were repeated in the presence of cumulatively rising concentrations of test compound applied by continuous perfusion, each concentration was present for 30 s before the recording started. The signals were sampled at 5 kHz after low-pass filtering at 2 kHz.

### Electrophysiological analysis

Data from automated and manual patch-clamp experiments were analyzed using Sophion Analyzer 10.0 software (Sophion Biosciences, Ballerup, Denmark) and Clampfit 10.7 (Molecular Devices, San Jose,CA), respectively, and GraphPad Prism 10 (version 10.0.2; GraphPad Software Inc., CA, USA). Ohmic leakage was subtracted from tail currents, and the leak-subtracted tail currents were plotted directly against the preceding test voltages to construct G(V) curves. These data were fit with a single Boltzmann curve:

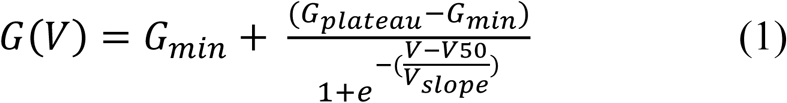

where G_min_ is the base, G_plateau_ is the top, V is the voltage in mV, V_50_ is the voltage where half of the maximum current is reached, and slope is the slope of the curve. To set the bottom of the curve to zero, the G_min_ was subtracted from the leak-subtracted tail currents, and data were again plotted directly against preceding test voltages and fit with a single Boltzmann curve:

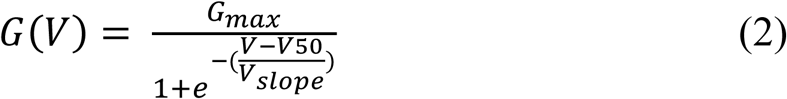

where G_max_ is the top, V is the voltage in mV, V_50_ is the voltage where half of the maximum current is reached, and slope is the slope of the curve. The slope was set to 8 and 12 for hERG and K_V_7.1/KCNE1, respectively. G_max_ and V_50_ values were exported from Sophion Analyzer (via Excel) for further analysis with GraphPad Prism.

Ratio of G_max_ (G/G_max_), and ΔV_50_ were calculated using the “Remove Baseline and Column Math” feature in GraphPad Prism 10, using the cell-specific 3^rd^ vehicle as the baseline, with ratio and difference calculation for G/G_max_ and ΔV_50_, respectively, and with sub-columns defined as repeated measures. G/G_max_ and ΔV_50_ were corrected for potential non-compound dependent effects in the recordings (i.e, run-down or run-up), again using the “Remove Baseline and Column Math” feature in GraphPad Prism 10, with ratio and difference calculation for G/G_max_ and ΔV_50_, respectively, using the corresponding data set for the run-specific vehicle as baseline, and with sub-columns defined as replicates. These normalized data, called G_max_ (rel.) and ΔV_50_ were then used for further analysis.

To determine concentration-dependence of responses to test compounds, G_max_ (rel.) and ΔV_50_ values were plotted and the following 4-parameter concentration-response curve was fitted to the data:

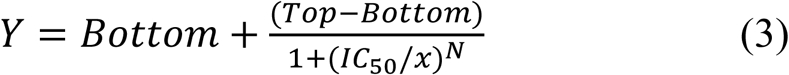

where *Y* is the response (G_max_ (rel.) or ΔV_50_), IC_50_ is the concentration that evokes 50% of the maximum response (constrained to always be above 0), and *N* is the Hill slope. For G_max_ (rel.), *Bottom* represents the maximally inhibited response and was constrained to assume a value between 0 and 1, and *Top* represents the baseline (no current inhibition) and was constrained to always be equal to 1. For ΔV_50_, *Bottom* represents the baseline (no shift in V_50_) and was constrained to always be equal to 0, while *Top* was the maximal V_50_ response and was not constrained.

### *In vitro* biological activity at CB_1_ receptor

The apoaequorin system (AequoScreen^®^ assay) is an intracellular calcium (Ca^2+^) release assay commonly used for measuring G-protein coupled (GPC) receptor activity. Apoaequorin is a protein that in the presence of its substrate coelenterazine converts to the aequorin photoprotein, which has three Ca^2+^ binding sites ^62^. Binding of an extracellular ligand to the GPC receptor leads to the activation of a universal G protein subunit Gα16, which in turn triggers a series of downstream events, such as the activation of the phospholipase C (PLC) enzyme, that stimulates the release of intracellular Ca^2+^ ^62^. The Ca^2+^ then binds to the aequorin photoprotein, which oxidizes the coelenterazine leading to the release of photons that can be measured via a luminescence reader ^63^.

CHO-K1 cells stably expressing CB_1_, the apoaequorin enzyme, and the Gα16 subunit were maintained in a humidified atmosphere at 37°C and 5% CO_2_ in Ham’s F12 medium supplemented with 10% heat-inactivated FBS. To perform the assays, cells were trypsinized (10 min, 37°C), centrifuged (at 200 × *g*, 5 min, room temperature), counted, resuspended at 3 × 10^5^ cells/mL in DMEM/Ham’s F12 without phenol red, and supplemented with 15 mM HEPES, L-glutamine, and protease-free BSA (0.1%) (further referred to as assay medium).

The coelenterazine substrate was added to a final concentration of 2.5 μM and the suspension was incubated for 3 h (room temperature, rotating at ∼7 RPM/min, protected from light). Drug solutions were prepared as a 1:8 serial dilution in assay medium (1:4 serial dilution for Δ^9^-THC) with a starting concentration of 60 μM (in well after addition of cells) and then added to white, opaque-welled 96-well plates. JWH-018 (60 μM) was included as a reference on each plate, in line with earlier results generated using the AequoScreen^®^ CB_1_ assay ^64–66^. Digitonin (67 μM) and ATP (6.7 μM) were included on each plate and served as positive controls for coelenterazine loading as both are involved in the non-CB-dependent release of calcium ions. Blank assay medium was used as a negative control. Using a TECAN Spark 10 M plate reader (Männedorf, Switzerland), 50 μL of the incubated cell suspension was dispensed into each well (15 × 10^3^ cells/well) of the 96-well plate containing the test solutions. Luminescence was measured for 25 s (corresponding to 190 additional reading cycles).

Absolute luminescence signals were corrected for intra-plate variability in Microsoft Excel 365 using area under the curve (AUC) values of JWH-018 and were calculated for each concentration of the test compounds. Values were then blank-corrected by subtracting AUC values of the mean of the blank controls. Data were normalized to the maximum of JWH-018. The normalized values were transferred to GraphPad Prism (Version 10.0.2) to generate concentration-response curves and calculate EC_50_ and E_max_ values by curve fitting via nonlinear regression (three-parameter logistic fit). Results are represented as receptor activity of JWH-018 (%) derived from a minimum of three independent experiments (n ≥ 3), run in triplicate. There were only two independent experiments (n = 2) for JWH-030, as there was not enough of the reference standard available for a third independent experiment. As the CB_1_ receptor activity of the SCRAs included in this study were examined over the span of a couple years, EC_50_ values were normalized to the median EC_50_ value of JWH-018 (35.03 nM) from 139 independent experiments run over many years.

### SAR-studies

Structure-activity relationship (SAR) analysis for hERG was conducted both for efficacy (maximum reduction in G_max_) and potency (IC_50_). Only comparisons involving at least three pairs of structurally related compounds were included. Compounds were only included if a concentration–response curve could be fitted; compounds with no measurable effect were excluded. SAR analysis for the CB_1_ receptor was conducted for both efficacy (E_max_) and potency (EC_50_), where the logEC_50_ values were used for the statistical analysis. Only comparisons involving at least three pairs of structurally related compounds were included.

### Statistical analysis

Data are reported as mean ± SEM. GraphPad Prism 10 was used for all statistical analyses. For comparisons between compound and run-specific vehicle controls in APC, a two-way ANOVA (Gaussian distribution of residuals and equal SDs assumed) followed by Dunnett’s multiple comparisons test was used. When a compound had been tested in several runs, the ANOVA (or mixed effect model when appropriate) was instead performed on the combined dataset and compared to the combined vehicle controls from corresponding runs, where all data were normalized according to their run prior to pooling. This was followed by either Dunnett’s multiple comparisons test (if more than one compound was to be compared to vehicle control) or a Šidák correction (if only one compound was to be compared to vehicle control). Correlation analysis was done with simple linear regression, accounting for n and scatter among replicates. Correlation was considered significant if the P-value of the slope significantly deviated from 0. For SAR studies on hERG, compounds were only included if a concentration–response curve could be fitted; compounds with no measurable effect were excluded. Log_10_IC_50_ values were used for statistical analysis. P values for compound-to-compound comparisons were obtained using extra-sum-of-squares F-tests and linear regression. P values for overall comparisons were calculated using two-tailed paired t-tests. A p-value < 0.05 was considered statistically significant. For SAR studies on CB_1_, P values for compound-to-compound comparisons were obtained using Brown-Forsythe and Welch ANOVA tests (α = 0.05) performed in GraphPad Prism, while p values for overall comparisons were calculated using two-tailed paired t-tests (α = 0.05).

### Molecular docking

To determine an initial binding pose for astemizole, the hERG channel structure (PDB ID: 7CN1, ^32^) was obtained from the RCSB protein data bank. Autodock Tools was utilized to prepare structures of protein and ligand for docking. Autodock Tools was also used to select a grid box of 24 Å centred on residue Y652 in the vestibule of the hERG channel. Autodock Vina ^67^ was utilized to calculate docking of astemizole within the box. 20 independent docking runs were attempted for each ligand and 15 poses were generated for each run. Outputs were clustered according to the root mean square deviation (RMSD), and the initial binding pose was used to guide the accelerated weighted histogram simulations.

### System set-up for molecular dynamics simulations

The inactivated-state hERG channel with astemizole bound (PDB entry: 7CN1, ^32^) from residue T399-L666 was embedded in a 100% POPC bilayer and solvated in 0.15 KCl and TIP3P water using CHARMM-GUI Membrane Builder. A single ligand (astemizole or SCRAs) was aligned with the initial binding pose of astemizole (determined by docking) using PyMOL to generate an initial configuration. Ligand parameters were generated using Ligand Reader and Modeler on CHARMM-GUI. The system was then energy-minimized and equilibrated using the standard six steps CHARMM-GUI equilibration protocol ^68^. This includes the following set-up: protein backbone was restrained at the force constant of 4000, 2000, 1000, 500, 200 and 50 kJmol^−1^ nm^−2^, protein side chains were restrained at the force constant of 2000, 1000, 500, 200, 50, and 0 kJmol^−1^ nm^−2^, non-H atoms of lipids were restrained at the force constant of 1000, 400, 400, 200, 40 and 0 kJmol^−1^ nm^−2^, and dihedral restraints were set at the force constant of 1000, 400, 200, 200, 100 and 0 kJmol^−1^ rad^−2^. Simulations were equilibrated with a 1 fs timestep for 125 ps for the first three steps, and then to a 2 fs timesteps for 500 ps the next two, and 5 ns for the final step. The first two steps were conducted with the NVT ensemble and the last four were conducted with the NPT ensemble. All equilibration runs were conducted at 310 K using a Berendsen thermostat ^69^. In all NPT ensemble equilibration, the pressure was maintained at 1 bar using a Berendsen barostat ^69^. All simulations were conducted using GROMACS-2023.4 ^70^. All images from simulations were generated using PyMOL.

### Accelerated weighted histogram simulations

The free energy landscape describing the path (z-axis distance) of the drug entering the binding site within the hERG channel was calculated using an accelerated weighted histogram (AWH) ^71^. Pressure was maintained at 1 bar and temperature was maintained at 310 K. An independent AWH bias was applied for each equilibrated structure, simulating 4 walkers for 1250 ns per walker, sharing bias data and target distribution. Bias acted on the z-axis, defined using the centre-of-mass of the drug and the Y652 residue in the vestibule of the hERG channel, aiming for a converged simulation, defined as a similar free energy landscape every 50 ns. Sampling interval was 1.0 nm above Y652 and 3.5 nm below Y652. The system initialized with an average free energy error of 20 kJ/mol, with the diffusion coefficient at 0.0002 nm^2^/ps at a force constant of 12800 kJ mol^−1^ nm^−2^. A harmonic potential was applied on all Cα atoms at 5.0 kJ mol^−1^ nm^−2^. Free energy landscape from 1210, 1220, 1230, 1240 and 1250 ns were averaged and used to build a final free energy landscape. The binding pose was clustered using DBSCAN with epsilon of 0.2 and minimum sample of 10. From each cluster, 1000 structures located closest to the free energy minima were extracted and used for minimum distance calculation.

## Acknowledgement

We acknowledge Dr. Fredrik Elinder, Linköping University, for valuable input on the study. The hERG Y652A clone was a generous gift from Dr. Nicole Schmitt, University of Copenhagen. This study was supported by funding from the European Research Council (ERC) under the European Union’s Horizon 2020 research and innovation program (grant agreement No. 850622), the Swedish Research Council (2021-01885), and the Strategic Research Area in Forensic Science, Linköping University. T.P. was funded by Lee Kuan Yew Postdoctoral Fellowship. C.N. was funded by a Postdoctoral Fellowship from the Swedish Society for Medical Research. Computational resources were supported by NTU-HPC center. The authors also acknowledge the support from the Chemical Biology Consortium Sweden (CBCS), node Linköping University, Electrophysiology, a national research infrastructure funded by the Swedish Research Council (2021-00179) and SciLifeLab. We acknowledge B’SYS GmbH for their collaboration and support in providing ion channel-expressing cell lines. We also thank Sophion Bioscience for their contributions to assay development, with special appreciation to Naja Møller Sørensen, Anissa Bara, and Rasmus Jacobsen for their expert assistance and valuable input throughout the project.

## Author contributions

Conceptualization: N.E.O., D.J.A.F, T.P., A.S., H.G., S.I.L.; Investigation and formal analysis: N.E.O., D.J.A.F, C.N., U.K., A.J.-M., M.V. for experimental work and T.P., A.S. for computational work; Supervision: N.E.O., H.P.L., H.G., S.I.L.; Writing – original draft: the original draft was mainly written by N.E.O., D.J.A.F, T.P., C.N., S.I.L. All authors contributed to and commented on the original draft.

## Declaration of interests

The authors declare no competing interests.

## Supplementary Figures

**Supplementary Figure S1.**
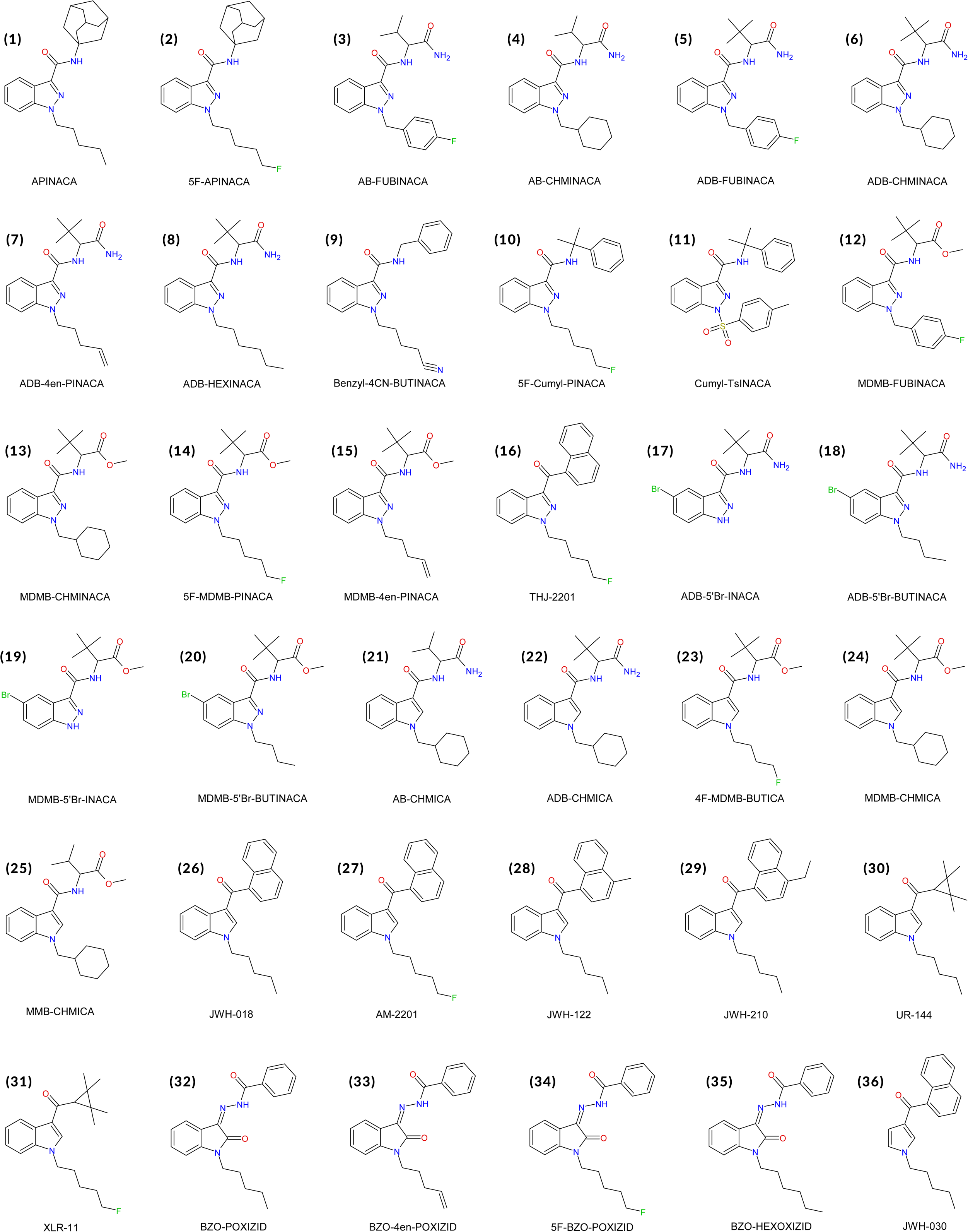
Molecular structures of the SCRAs included in this study. SCRAs are numbered according to the list order in Supplementary Table I.

**Supplementary Figure S2:**
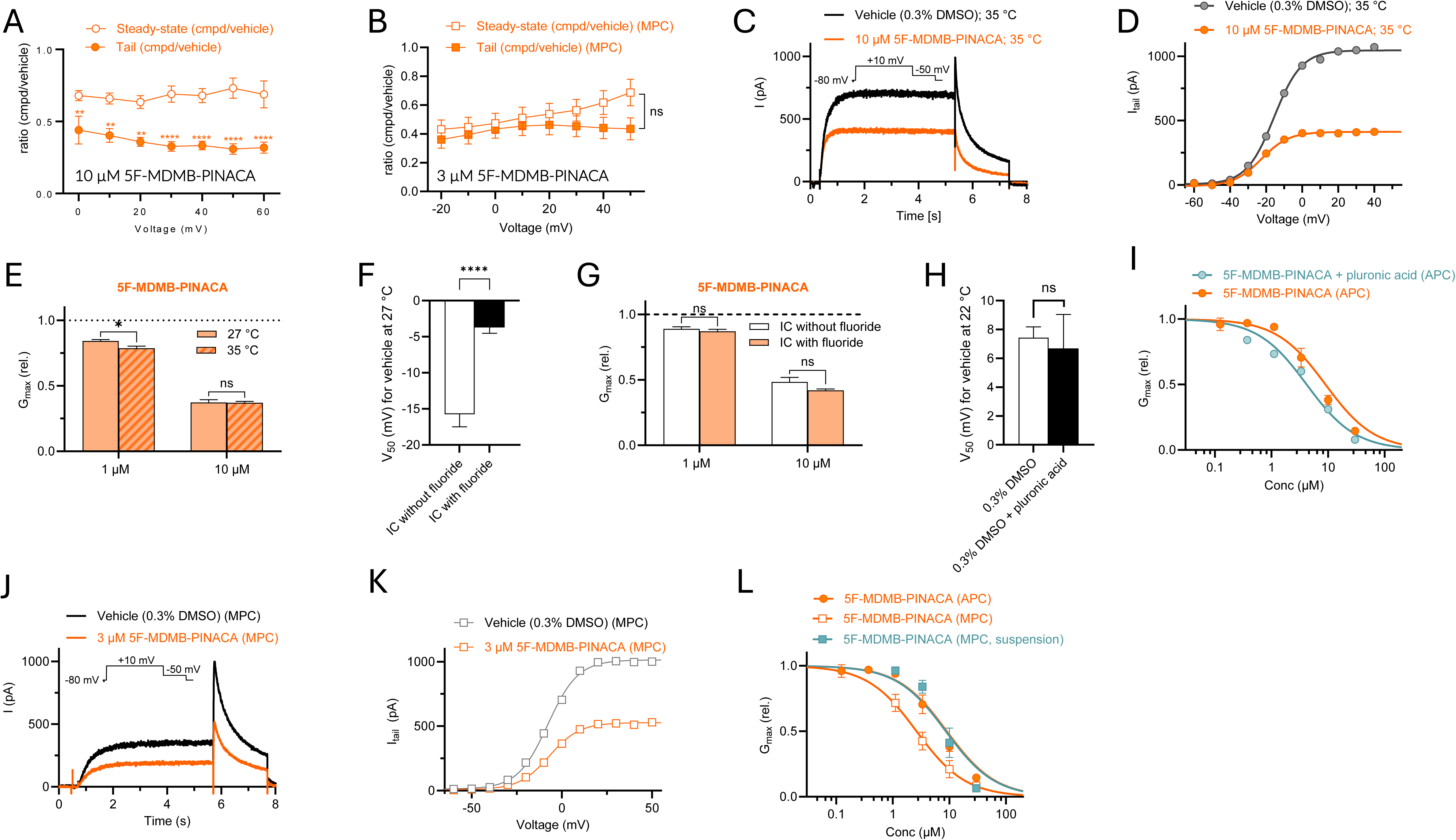
Electrophysiological characterization of 5F-MDMB-PINACA inhibition on wild type (WT) hERG highlight aspects impacting effects. (A-B) Average ratio of 5F-MDMB-PINACA effects on steady-state current compared to tail current, collected using 10 µM 5F-MDMB-PINACA for APC (A) and 3 µM 5F-MDMB-PINACA for MPC (B). Statistics denote difference between effects on steady-state current compared to tail current, compared using two-way ANOVA together with Šídák’s multiple comparisons test, p < 0.05 (*). n = 5-9 for APC, n = 13 for MPC. **(C-D)** Representative effect of 10 µM 5F-MDMB-PINACA on hERG at 35°C, collected using APC and the pulse protocol shown as insert (C), with corresponding G(V) curve (D). **(E)** Average effect of indicated concentrations of 5F-MDMB-PINACA on hERG at 27°C or 35°C, collected using APC. Data are presented as mean ± SEM. Statistical comparisons indicate differences between temperatures at a given concentration, mixed-effect analysis with Šídák’s multiple comparisons test, (p < 0.05), n = 3–9. **(F)** Average voltage-dependence of activation (V_50_) for hERG at 27°C with or without fluoride in the intracellular solution. Data are presented as mean ± SEM. Statistical comparisons indicate difference between with or without fluoride in the intracellular solution, unpaired t-test, p < 0.001 (***), n = 12-13. **(G)** Average effect of indicated concentrations of 5F-MDMB-PINACA on hERG at 27°C, with or without fluoride in the intracellular solution, collected using APC. Data are presented as mean ± SEM. Statistical comparisons indicate differences between with or without fluoride in the intracellular solution, at a given concentration, mixed-effect analysis with Šídák’s multiple comparisons test, (p > 0.05), n = 7–11. (**H**) Average voltage-dependence of activation (V_50_) for hERG at 22°C with or without pluronic acid in the vehicle. Data are presented as mean ± SEM. Statistical comparisons indicate difference between with or without pluronic acid in the vehicle, unpaired t-test, p > 0.05, n = 5. (**I**) Concentration-response relationship for the G_max_ (rel) effect of 5F-MDMB-PINACA on hERG without pluronic acid (orange) and with pluronic acid (teal), collected using APC. Best fits for G_max_ (rel) yielded IC_50_ values of 9.0 µM (n = 4-17) without pluronic acid and 4.2 µM (n = 10-11) with pluronic acid. The IC_50_ values were significantly different (p < 0.0001) using Extra sum-of-squares F-test. For both with and without pluronic acid, the maximal inhibition approached full inhibition. (**J-K**) Representative effect of 3 µM 5F-MDMB-PINACA on hERG at 22°C, collected using MPC and the pulse protocol shown as insert (J), with corresponding G(V) curve (K). (**L**) Concentration-response relationship for the G_max_ (rel) effect of 5F-MDMB-PINACA on hERG collected using APC with cells in suspension (orange, filled circles); using MPC with attached cells (orange, unfilled squares); and using MPC with cells in suspension (teal, filled squares). Best fits for G_max_ (rel) yielded IC_50_ values of 9.0 µM (n = 4-17) for APC with cells in suspension; 2.6 µM (n = 5-13) for MPC with attached cells; and 8.6 µM (n = 2-3) for MPC with cells in suspension. The IC_50_ values for cells in suspension recorded using APC and attached cells recorded using MPC were significantly different (p = 0.0001). For cells in suspension, the IC_50_ values for recordings using APC and MPC were not significantly different (p = 0.08). For recordings using MPC, the IC_50_ values for attached cells and cells in suspension were significantly different (p = 0.0093). For all three, the maximal inhibition approached full inhibition. Statistical calculations were done using Extra sum-of-squares F-test.

**Supplementary Figure S3.**
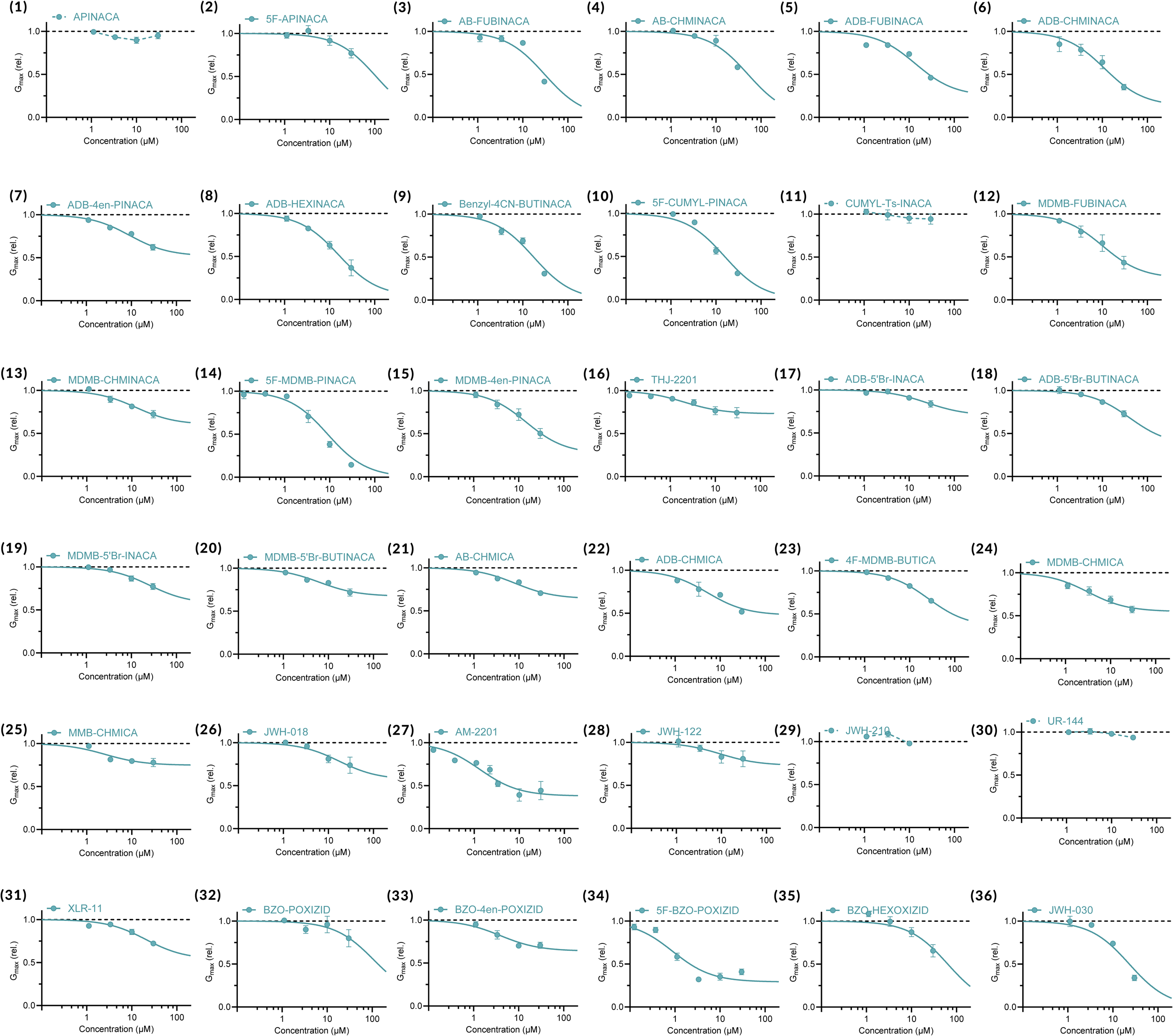
Concentration-response relationships for SCRA effects on G_max_ of hERG. All average data are shown as mean ± SEM. Small error bars are covered by symbols. n = 3-13. Best fits as in SI Table II. For SCRAs with no clear inhibitory effects, data points are connected with dashed lines. SCRA numbering is according to Supplementary Figure S1 and Supplementary Table I.

**Supplementary Figure S4.**
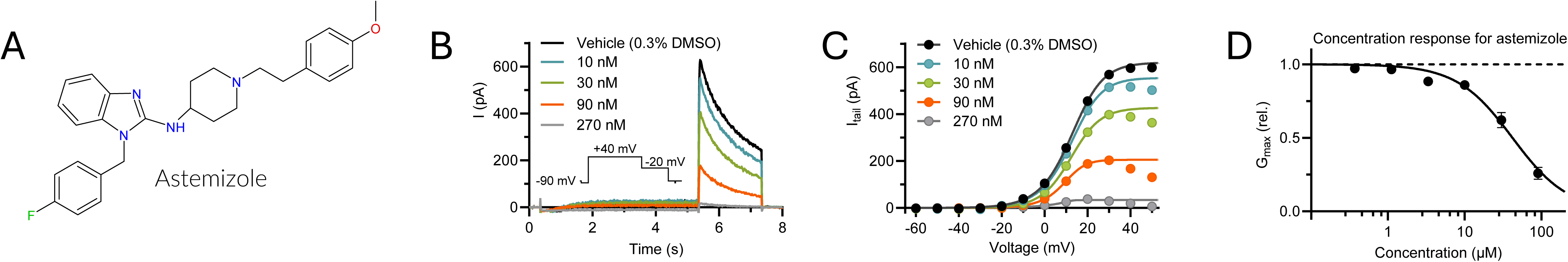
Effect of the hERG blocker astemizole in our APC hERG assay. **(A)** Molecular structure of astemizole. **(B-C)** Representative effect of astemizole on hERG using the pulse protocol shown as insert in B, with corresponding G(V) curves in C. **(D)** Concentration-response relationships for astemizole on hERG inhibition. Best fit indicates IC_50_ = 41.9 ± 9.2 nM. n = 4. Data shown as mean ± SEM. Small error bars are covered by symbols.

**Supplementary Figure S5.**
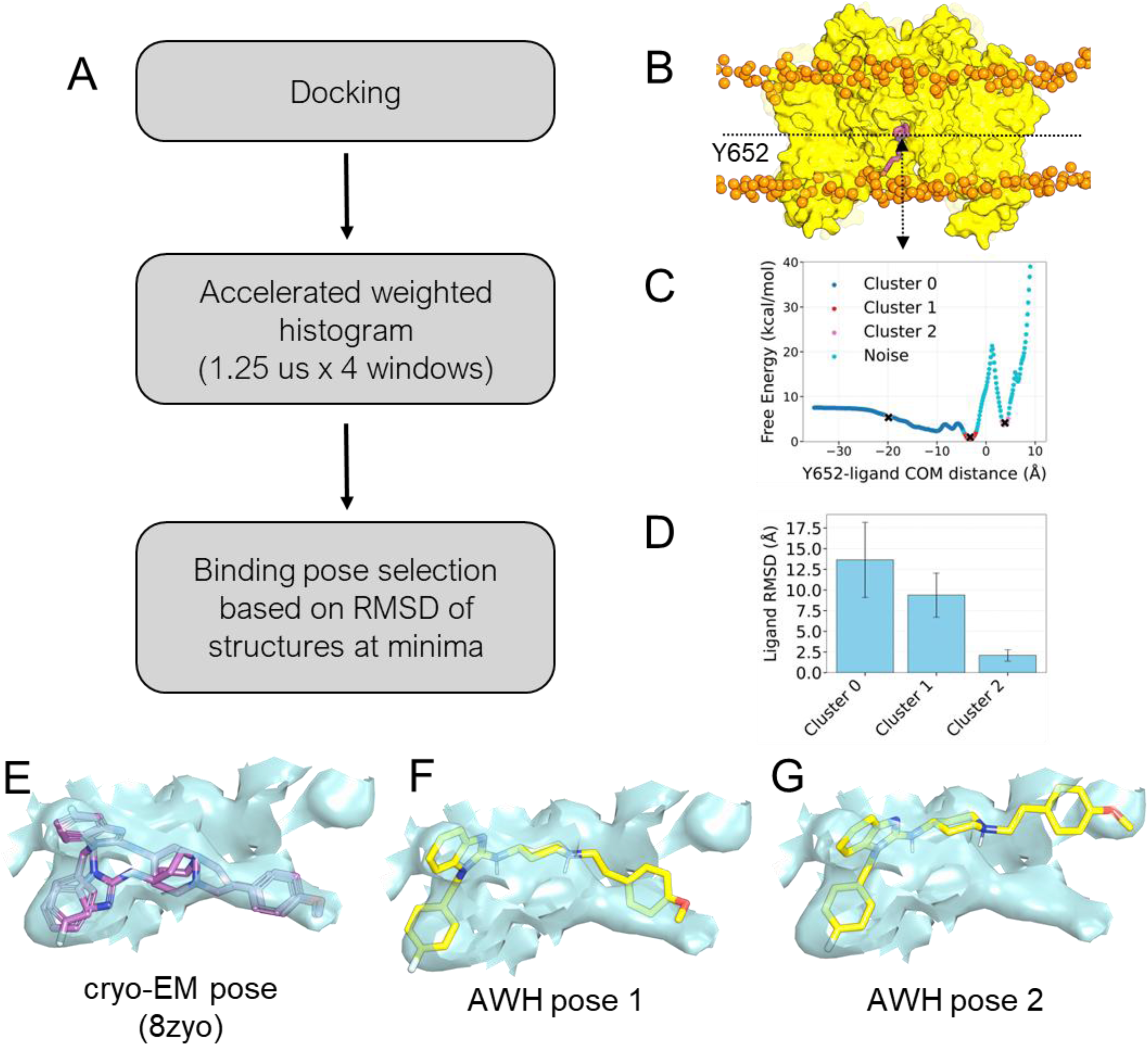
AWH workflow used to determine SCRA binding. **(A)** *In silico* workflow employed to obtain the binding pose of astemizole and SCRAs. **(B)** The structural representation of the sampling of the astemizole ligand along the z-axis with respect of Y652. (C) Free energy landscape of astemizole from AWH simulations using 4 windows simulating for 1250 ns. The region highlighted with the cross shows the energy minima from each cluster. **(D)** Root mean square deviation of the ligand within the cluster, calculated from 1000 ligands in the binding site calculated with respect to the cluster centre. The error bar shows the standard deviation of the RMSD. **(E-G)** The electron density map (cyan) of the ligand binding site, fitting around the ligand astemizole. The cryo-EM model (PDB entry: 8ZYO) is shown in purple (E). The frames from Cluster 1 from AWH simulations in two distinct conformations are shown in yellow (F-G).

**Supplementary Figure S6.**
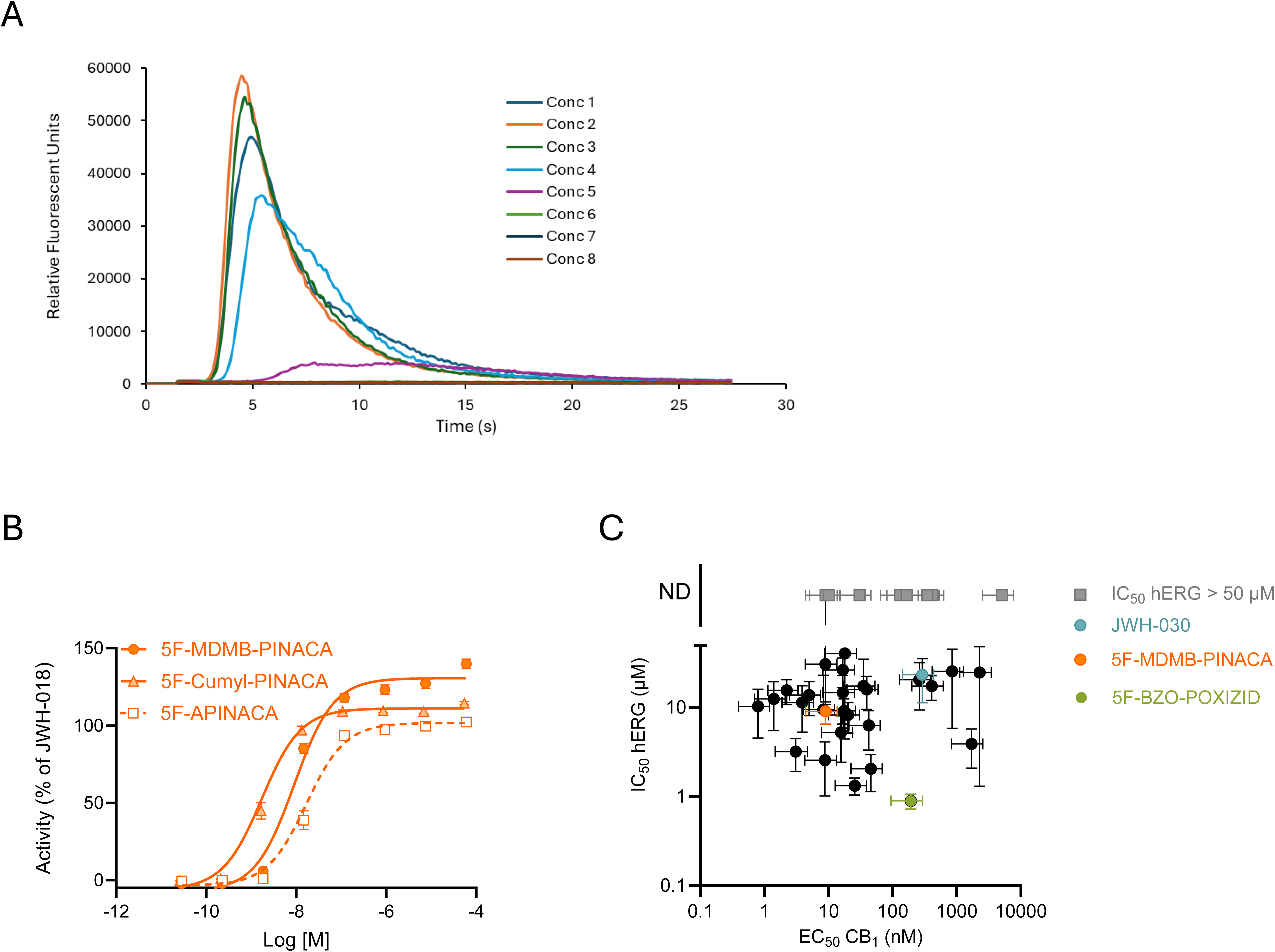
CB_1_ receptor activity assay. **(A)** Representative effect of JWH-018 on CB_1_ receptor activity. The fluorescence signal was recorded over time of 8 concentrations of JWH-018 tested on the AequoScreen® CB_1_ receptor assay. **(B)** Example of impact of head chemistry for CB_1_ activation showcased by 5F-PINACA compounds with either MDMB, Cumyl, or adamantyl heads. Data is from a minimum of three independent experiments (n ≥ 3), run in triplicate. **(C)** Comparison of SCRA potency for hERG and CB_1_. Specific SCRAs are highlighted as indicated in the figure. Note that SCRAs marked in grey were either non-inhibitors of hERG or had an anticipated IC_50_ exceeding 50 µM. All average data are shown as mean ± SEM. Small error bars are covered by symbols. Best fits as in SI Table II.

**Supplementary Figure S7.**
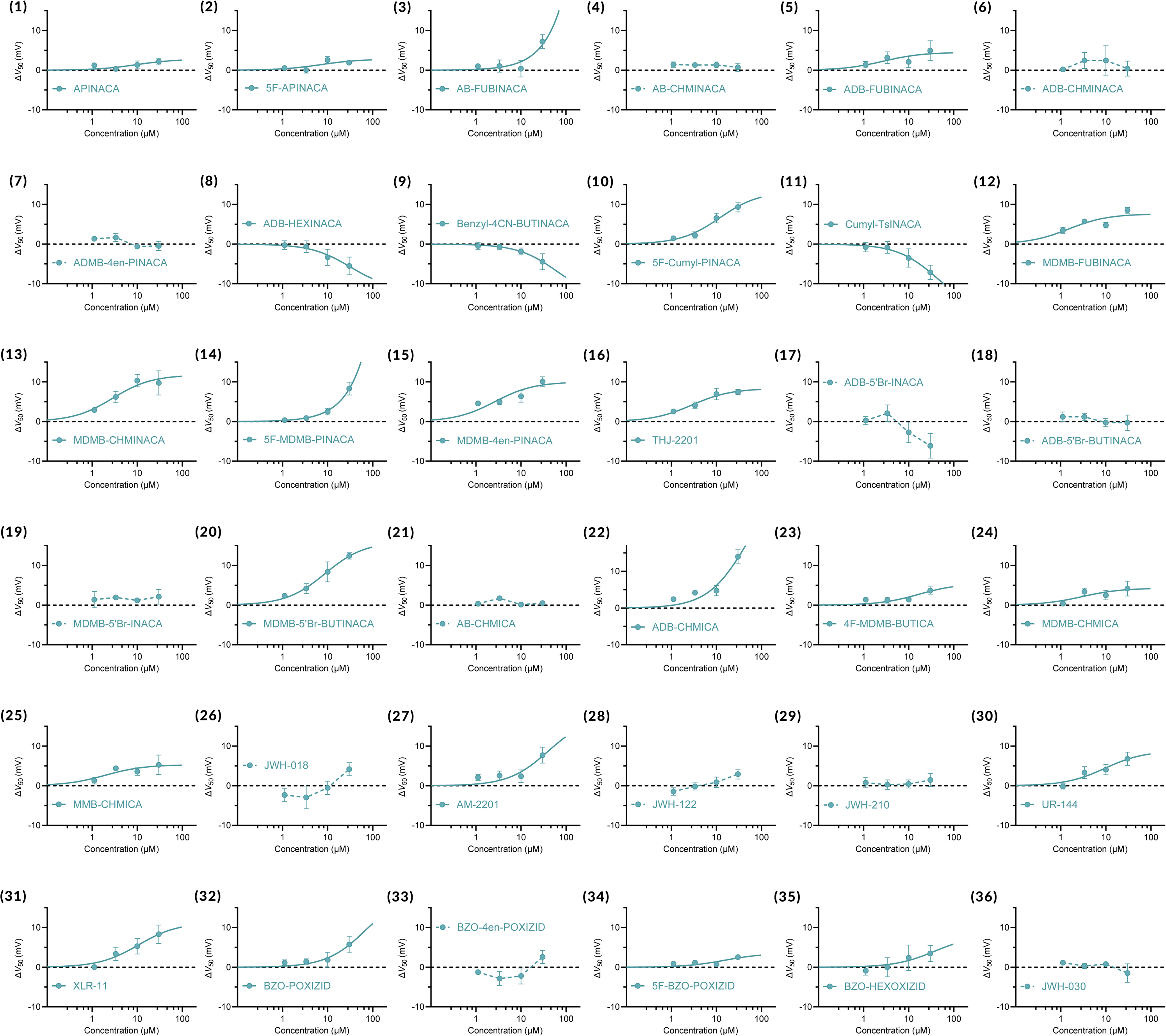
Concentration-response relationships for SCRA effects on V_50_ of K_V_7.1/KCNE1. All average data are shown as mean ± SEM. Small error bars are covered by symbols. n = 3-9. For SCRAs with no clear V_50_ effects, data points are connected with dashed lines. SCRA numbering is according to Supplementary Figure S1 and Supplementary Table I.

**Supplementary Figure S8.**
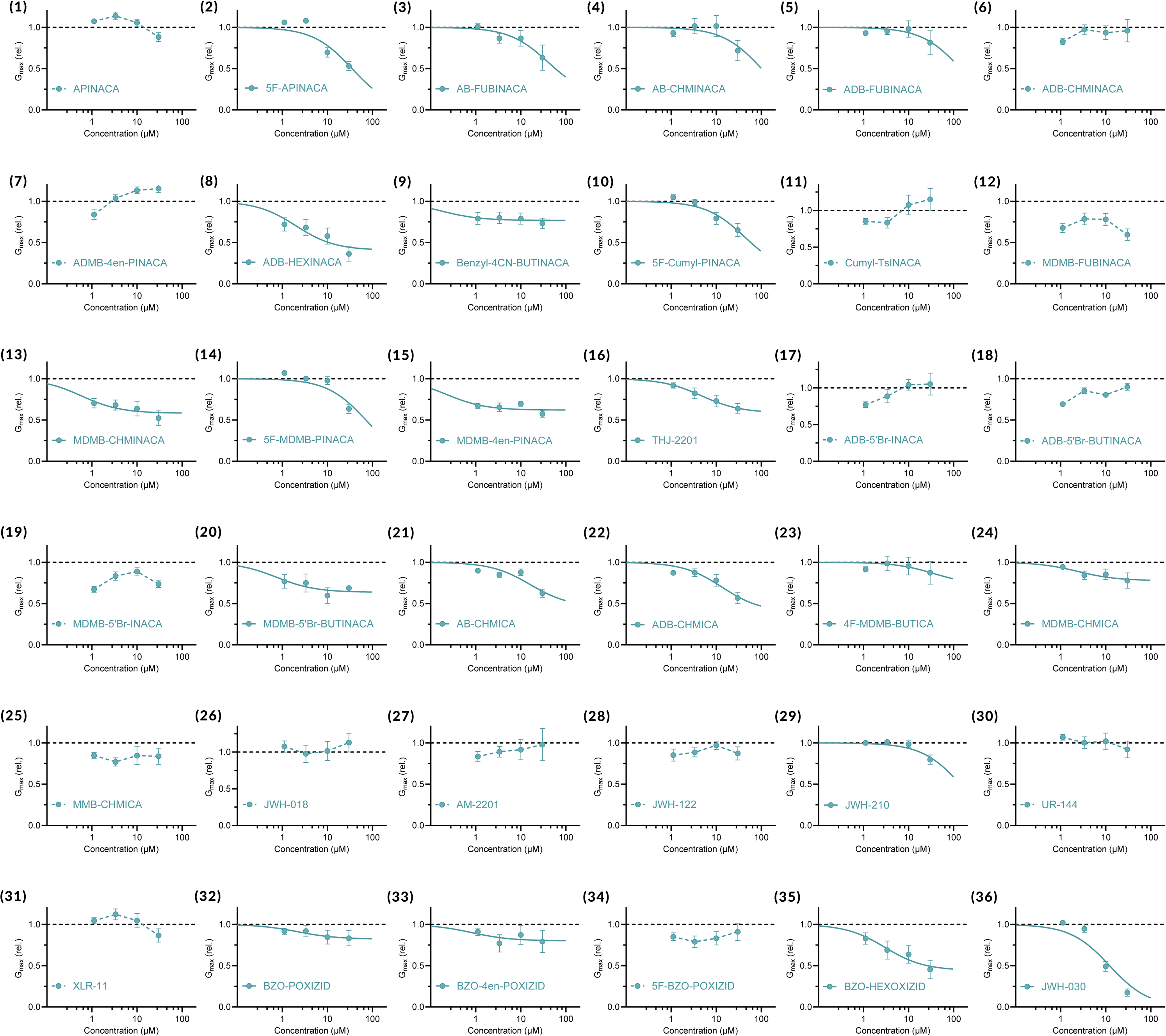
Concentration-response relationships for SCRA effects on G_max_ of K_V_7.1/KCNE1. All average data are shown as mean ± SEM. Small error bars are covered by symbols. n = 3-9. Best fits as in SI Table II. For SCRAs with no clear G_max_ effects, data points are connected with dashed lines. SCRA numbering is according to Supplementary Figure S1 and Supplementary Table I.

## Supplementary Text

### Experimental characterization of 5F-MDMB-PINACA effects on hERG, to explore experimental conditions potentially impacting inhibitory effects

To more carefully assess the inhibitory effects of the SCRAs on hERG channel function, we performed additional analysis and experiments using 5F-MDMB-PINACA, which was one of the most prominent hERG inhibitors in our APC assay. For APC recordings of 5F-MDMB-PINACA, we noted a more pronounced reduction of the tail current compared to the steady-state current across voltages with a clear channel activation (SI Fig. S2A). This difference was not present in the corresponding MPC recordings (SI Fig. S2B). In both methods, the unspecific ohmic leak is compensated for using the voltage step prior to activation. In part, challenges in fully compensating for potassium ion leak in APC recordings may have contributed to the relatively larger tail current effects. However, additional factors such as the time course of the onset of 5F-MDMB-PINACA effects and possibly some degree of state dependence of 5F-MDMB-PINACA effects may contribute.

Using APC, we examined the effect of temperature on SCRA activity by performing recordings at 27°C and 35°C. While increased temperature accelerated channel kinetics (Fig. S2C), the inhibitory effect of 10 µM 5F-MDMB-PINACA was unaffected by the increased temperature (SI Fig. S2D-E) whilst the inhibitory effect of 1 µM 5F-MDMB-PINACA was slightly larger at the higher temperature (Fig. S2E). The intracellular solution used in our APC and MPC recordings contained fluoride (24 mM) to improve seal quality and recording stability ^72^. We observed a shift in the voltage dependence of channel activation in vehicle control experiments in the presence of fluoride by +12 mV (SI Fig. S2F, recordings were done at 27 °C). However, the inhibitory effects of 5F-MDMB-PINACA on both steady-state currents and tail currents were unaffected by the presence of fluoride (SI Fig. S2G). These data suggest that 5F-MDMB-PINACA acts independently of fluoride and its inhibitory effect is relatively temperature-independent across a range of temperatures.

Due to their high lipophilicity, SCRAs such as 5F-MDMB-PINACA may interact with hydrophobic components of the recording environment, including surfaces or materials within the APC system, even in the absence of plastic. This property could influence compound availability and apparent potency. To minimize this, pluronic acid was included as a carrier in specific experiments. Pluronic acid did not affect the voltage dependence of channel activation (SI Fig. S2H) but enhanced compound potency, as reflected by a lower IC_50_ for 5F-MDMB-PINACA (SI Fig. S2I). Manual patch-clamp (MPC) recordings revealed an even greater potency, with 3 µM producing a comparable effect to 10 µM in APC (SI Fig. S2J-K). A major methodological difference between MPC and APC is that the MPC recordings were performed on single, adherent cells, whereas APC recordings were performed on cells that had been enzymatically detached from the culture surface using Detachin™ and kept in suspension during measurements. To investigate if cell detachment influenced compound potency, we also performed MPC recordings on suspended cells.

Interestingly, the IC_50_ for 5F-MDMB-PINACA was higher for detached cells compared to adherent ones (SI Fig. S2L), approaching the IC_50_ for 5F-MDMB-PINACA in APC. Consequently, the potency of SCRAs on hERG may be greater than what is indicated by APC recordings using suspended cells, underscoring a role of cell state in pharmacological assessments.

---

**Supplementary Table I:** The 36 SCRAs included in this study organized by their four main structural components (core, linker, head group, and tail). For schematic example, see Figure 1C. Molecular structures of all SCRAs are shown in Supplementary Figure S1.

|  | SCRA name | Other used name(s) | Core | Linker | Head | Tail |
| --- | --- | --- | --- | --- | --- | --- |
| 1 | APINACA | A-PINACA, AKB-48 | Indazole | Carboxamide | Adamantyl | Pentyl |
| 2 | 5F-APINACA | A-5F-PINACA, 5F-AKB-48 | Indazole | Carboxamide | Adamantyl | 5-fluoropentyl |
| 3 | AB-FUBINACA | AMB-FUBINACA | Indazole | Carboxamide | AB | 4-fluorobenzyl |
| 4 | AB-CHMINACA | AMB-CHMINACA | Indazole | Carboxamide | AB | Cyclohexylmethyl |
| 5 | ADB-FUBINACA | ADMB-FUBINACA | Indazole | Carboxamide | ADB | 4-fluorobenzyl |
| 6 | ADB-CHMINACA | ADMB-CHMINACA | Indazole | Carboxamide | ADB | Cyclohexylmethyl |
| 7 | ADB-4en-PINACA | ADMB-4en-PINACA | Indazole | Carboxamide | ADB | Pent-4-enyl |
| 8 | ADB-HEXINACA | ADMB-HINACA | Indazole | Carboxamide | ADB | Hexyl |
| 9 | Benzyl-4CN-BUTINACA | BZ-4CN-BINACA | Indazole | Carboxamide | Benzyl | 4-cyanobutyl |
| 10 | 5F-Cumyl-PINACA | CUMYL-5F-PINACA | Indazole | Carboxamide | Cumyl | 5-fluoropentyl |
| 11 | Cumyl-TsINACA | CUMYL-TSINACA | Indazole | Carboxamide | Cumyl | Tosyl |
| 12 | MDMB-FUBINACA |  | Indazole | Carboxamide | MDMB | 4-fluorobenzyl |
| 13 | MDMB-CHMINACA |  | Indazole | Carboxamide | MDMB | Cyclohexylmethyl |
| 14 | 5F-MDMB-PINACA | MDMB-5F-PINACA, 5F-ADB | Indazole | Carboxamide | MDMB | 5-fluoropentyl |
| 15 | MDMB-4en-PINACA |  | Indazole | Carboxamide | MDMB | Pent-4-enyl |
| 16 | THJ-2201 | NA-5F-PINAMO | Indazole | Methanone | 1-naphthyl | 5-fluoropentyl |
| 17 | ADB-5'Br-INACA | ADMB-5Br-INACA | 5-bromo-1H-indazole | Carboxamide | ADB | - |
| 18 | ADB-5'Br-BUTINACA | ADMB-B-5Br-INACA | 5-bromo-1H-indazole | Carboxamide | ADB | Butyl |
| 19 | MDMB-5'Br-INACA | MDMB-5Br-INACA | 5-bromo-1H-indazole | Carboxamide | MDMB | - |
| 20 | MDMB-5'Br-BUTINACA | MDMB-B-5Br-INACA | 5-bromo-1H-indazole | Carboxamide | MDMB | Butyl |
| 21 | AB-CHMICA | AMB-CHMICA | Indole | Carboxamide | AB | Cyclohexylmethyl |
| 22 | ADB-CHMICA | ADMB-CHMICA | Indole | Carboxamide | ADB | Cyclohexylmethyl |
| 23 | 4F-MDMB-BUTICA | MDMB-4F-BICA | Indole | Carboxamide | MDMB | 4-fluorobutyl |
| 24 | MDMB-CHMICA |  | Indole | Carboxamide | MDMB | Cyclohexylmethyl |
| 25 | MMB-CHMICA | AMB-CHMICA | Indole | Carboxamide | MMB | Cyclohexylmethyl |
| 26 | JWH-018 | NA-PIMO | Indole | Methanone | 1-naphthyl | Pentyl |
| 27 | AM-2201 | NA-5F-PIMO, 5F-JWH-018 | Indole | Methanone | 1-naphthyl | 5-fluoropentyl |
| 28 | JWH-122 | 4Me-NA-PIMO | Indole | Methanone | 4-methyl-naphthalen-1-yl | Pentyl |
| 29 | JWH-210 | 4Et-NA-PIMO | Indole | Methanone | 4-ethyl-naphthalen-1-yl | Pentyl |
| 30 | UR-144 | TMCP-PIMO | Indole | Methanone | Tetramethylcyclopropyl | Pentyl |
| 31 | XLR-11 | TMCP-5F-PIMO, 5F-UR-144 | Indole | Methanone | Tetramethylcyclopropyl | 5-fluoropentyl |
| 32 | BZO-POXIZID | 5C-MDA-19 | 2-oxindole | Hydrazide | Benzoyl | Pentyl |
| 33 | BZO-4en-POXIZID |  | 2-oxindole | Hydrazide | Benzoyl | Pent-4-enyl |
| 34 | 5F-BZO-POXIZID | BZO-5F-POXIZID, 5F-MDA-19 | 2-oxindole | Hydrazide | Benzoyl | 5-fluoropentyl |
| 35 | BZO-HEXOXIZID | BZO-HOXIZID, MDA-19 | 2-oxindole | Hydrazide | Benzoyl | Hexyl |
| 36 | JWH-030 | NA-PPYOMO | 1H-pyrrole | Methanone | 1-naphthyl | Pentyl |

\*Abbreviations: AB = 1-amino-3-methyl-1-oxobutane; ADB = 1-amino-3,3-dimethyl-1-oxobutane; MDMB = methyl-3,3-dimethyl-butanoate; MMB = methyl-3-methyl-butanoate

**Supplementary Table II:** Efficacy (maximal reduction in Gmax) and potency (IC50) for the inhibition of the hERG channel and relative efficacy (Emax) and potency (EC50) for the activation of the CB1 receptor (where JWH-018 was used as the reference), calculated for the 36 SCRAs included in this study. SCRAs are organized by their four main structural components, as shown in Supplementary Table I.

| SCRA | hERG channel | | | | $CB_1$ receptor | | | | Kv7.1/KCNE1 channel | | | |
| --- | --- | --- | --- | --- | --- | --- | --- | --- | --- | --- | --- | --- |
| | Efficacy (max reduction in $G_{\max}$) | | Potency ($\mu M$) | | Efficacy (% JWH-018) | | Potency (nM) | | Efficacy (max reduction in $G_{\max}$) | | Potency ($\mu M$) | |
| | $G_{\min}$ | SEM | $IC_{50}$ | SEM | $E_{\max}$ | SEM | $EC_{50}$ | SEM | $G_{\min}$ | SEM | $IC_{50}$ | SEM |
| APINACA | 0.90* | ND | ND | ND | 108 | 1.05 | 30.4 | 15.5 | 0.88* | ND | ND | ND |
| 5F-APINACA | 0.00 | 3.45 | 106.7 | 458.2 | 102 | 1.22 | 16.6 | 8.5 | 0 | 0.63 | 35.11 | 36.7 |
| AB-FUBINACA | 0.00 | 0.37 | 30.4 | 19.6 | 108 | 1.30 | 8.94 | 4.6 | 0.15 | 1.11 | 40.84 | 81.57 |
| AB-CHMINACA | 0.00 | 0.72 | 51.2 | 54.3 | 108 | 1.49 | 8.98 | 4.6 | 0 | 0 | 101 | 783.43 |
| ADB-FUBINACA | 0.25 | 0.14 | 13.7 | 5.67 | 106 | 1.23 | 4.97 | 2.5 | 0 | 10.02 | 142.16 | 1690.99 |
| ADB-CHMINACA | 0.13 | 0.19 | 11.3 | 6.05 | 107 | 0.68 | 3.87 | 2.0 | 0.93* | ND | ND | ND |
| ADB-4en-PINACA | 0.52 | 0.067 | 9.10 | 3.27 | 117 | 1.77 | 17.4 | 8.9 | 1.13* | ND | ND | ND |
| ADB-HEXINACA | 0.037 | 0.19 | 15.8 | 6.35 | 117 | 1.21 | 39.4 | 20.1 | 0.41 | 0.11 | 1.99 | 1.31 |
| Benzyl-4CN-BUTINACA | 0.00 | 0.16 | 17.3 | 5.37 | 109 | 1.74 | 408 | 208.4 | 0.77 | 0.05 | 0.17 | 0.54 |
| 5F-Cumyl-PINACA | 0.00 | 0.15 | 15.5 | 4.82 | 112 | 1.13 | 2.23 | 1.1 | 0.13 | 0.88 | 43.45 | 68.06 |
| Cumyl-TsINACA | 0.94* | ND | ND | ND | 106 | 1.58 | 166 | 84.7 | 1.07* | ND | ND | ND |
| MDMB-FUBINACA | 0.25 | 0.16 | 10.2 | 5.68 | 103 | 1.26 | 0.795 | 0.4 | 0.59* | ND | ND | ND |
| MDMB-CHMINACA | 0.60 | 0.10 | 12.4 | 6.93 | 106 | 0.94 | 1.40 | 0.7 | 0.58 | 0.06 | 0.59 | 0.51 |
| 5F-MDMB-PINACA | 0.00 | 0.11 | 8.98 | 2.49 | 131 | 1.62 | 8.99 | 4.6 | 0 | 1.65 | 71.37 | 159.62 |
| MDMB-4en-PINACA | 0.27 | 0.18 | 14.6 | 8.13 | 109 | 1.06 | 16.7 | 8.5 | 0.62 | 0.03 | 0.22 | 0.24 |
| THJ-2201 | 0.73 | 0.037 | 2.04 | 0.91 | 92.8 | 1.17 | 45.6 | 23.3 | 0.92 | 0.08 | 0 | 2.24 |
| ADB-5'Br-INACA | 0.70 | 0.15 | 24.5 | 23.2 | 111 | 1.88 | 2290 | 1168.4 | 1.04* | ND | ND | ND |
| ADB-5'Br-BUTINACA | 0.36 | 0.36 | 40.1 | 35.7 | 110 | 1.20 | 18.0 | 9.2 | 0.80* | ND | ND | ND |
| MDMB-5'Br-INACA | 0.57 | 0.18 | 25.3 | 19.5 | 110 | 1.26 | 849 | 433.2 | 0.74* | ND | ND | ND |
| MDMB-5'Br-BUTINACA | 0.67 | 0.053 | 6.24 | 2.92 | 114 | 1.69 | 42.4 | 21.6 | 0.64 | 0.08 | 0.74 | 0.87 |
| AB-CHMICA | 0.64 | 0.051 | 8.18 | 3.08 | 110 | 0.92 | 20.1 | 10.2 | 0.46 | 0.18 | 16.19 | 11.31 |
| ADB-CHMICA | 0.48 | 0.10 | 5.22 | 2.80 | 110 | 0.41 | 15.7 | 8.0 | 0.4 | 0.19 | 13.41 | 9.56 |

| SCRA | hERG channel |  |  |  | CB<sub>1</sub> receptor |  |  |  | Kv7.1/KCNE1 channel |  |  |  |
| --- | --- | --- | --- | --- | --- | --- | --- | --- | --- | --- | --- | --- |
|  | Efficacy (max reduction in G<sub>max</sub>) |  | Potency (μM) |  | Efficacy (% JWH-018) |  | Potency (nM) |  | Efficacy (max reduction in G<sub>max</sub>) |  | Potency (μM) |  |
|  | G<sub>min</sub> | SEM | IC<sub>50</sub> | SEM | E<sub>max</sub> | SEM | EC<sub>50</sub> | SEM | G<sub>min</sub> | SEM | IC<sub>50</sub> | SEM |
| 4F-MDMB-BUTICA | 0.35 | 0.13 | 26.2 | 9.50 | 115 | 1.34 | 16.7 | 8.5 | 0.72 | 1.09 | 36.94 | 237.74 |
| MDMB-CHMICA | 0.55 | 0.051 | 3.18 | 1.28 | 107 | 0.69 | 3.07 | 1.6 | 0.78 | 0.07 | 2.4 | 2.9 |
| MMB-CHMICA | 0.75 | 0.041 | 2.55 | 1.54 | 109 | 0.91 | 8.76 | 4.5 | 0.83 | ND | ND | ND |
| JWH-018 | 0.57 | 0.21 | 17.3 | 17.3 | 100 | 0.41 | 35.0 | 17.9 | 1.01* | ND | ND | ND |
| AM-2201 | 0.38 | 0.042 | 1.32 | 0.282 | 99.7 | 1.62 | 25.8 | 13.1 | 0.92* | ND | ND | ND |
| JWH-122 | 0.73 | 0.16 | 9.39 | 13.5 | 105 | 1.34 | 8.29 | 4.2 | 0.87* | ND | ND | ND |
| JWH-210 | 0.89* | ND | ND | ND | 110 | 1.58 | 10.1 | 5.1 | 0 | 4.61 | 141.49 | 774.86 |
| UR-144 | 0.94* | ND | ND | ND | 101 | 1.73 | 414 | 211.4 | 0.92* | ND | ND | ND |
| XLR-11 | 0.54 | 0.13 | 20.4 | 11.1 | 105 | 0.96 | 260 | 132.7 | 0.87* | ND | ND | ND |
| BZO-POXIZID | 0.10 | 4.91 | 108 | 728 | 108 | 2.28 | 349 | 178.1 | 0.82 | 0.08 | 2.18 | 3.71 |
| BZO-4en-POXIZID | 0.64 | 0.054 | 3.88 | 1.81 | 112 | 3.32 | 1700 | 867.3 | 0.8 | 0.08 | 0.81 | 1.82 |
| 5F-BZO-POXIZID | 0.29 | 0.037 | 0.892 | 0.171 | 113 | 2.51 | 193 | 98.5 | 0.83* | ND | ND | ND |
| BZO-HEXOXIZID | 0.00 | 1.53 | 63.6 | 135 | 84.3 | 3.15 | 5130 | 2617.3 | 0.44 | 0.12 | 2.99 | 2.31 |
| JWH-030 | 0.00 | 0.28 | 23.1 | 11.9 | 76.5 | 0.80 | 291 | 148.3 | 0 | 0.2 | 12.16 | 5.74 |

Efficacy (maximal reduction in G<sub>max</sub>) is in the table indicated as G<sub>min</sub>, where 1 indicates no reduction of G<sub>max</sub> and 0 indicates complete reduction of G<sub>max</sub> (i.e., no current remaining). ND, not determined (values could not be calculated as saturation was not reached). \* Greatest efficacy achieved with a concentration between 10 and 30 μM. n = 3-13 for hERG and n = 3-9 for Kv7.1/KCNE1. For CB<sub>1</sub>, Results are represented as receptor activity of JWH-018 (%) derived from a minimum of three independent experiments (n ≥ 3), run in triplicate (except for JWH-030, for which there were only two independent experiments, n = 2).

**Supplementary Table III.** SAR analysis for impact of core moiety on hERG efficacy (maximal reduction in Gmax) and potency (IC50). P values for compound-to-compound comparisons are from Extra-sum-of-squares F-test and linear regression. P values for overall comparisons are from two-tailed paired t-tests. A negative Mean Diff indicates that Compound 1 has a bigger maximal effect (i.e., a larger maximal reduction in Gmax) and is more potent (i.e, has a smaller IC50), respectively.

| | Comparisons | | Efficacy (max reduction in $G_{\max}$) | | | log$IC_{50}$ | | |
| --- | --- | --- | --- | --- | --- | --- | --- | --- |
|  | Compound 1 | Compound 2 | Mean Diff | P value | Significant? | Mean Diff | P value | Significant? |
| **Indole**<br>**vs.**<br>**Indazole** | AB-CHMICA | AB-CHMINACA | 0.6460 | 0.0056 | Yes | -0.784 | 0.0099 | Yes |
|  | ADB-CHMICA | ADB-CHMINACA | 0.3439 | 0.3917 | No | -0.334 | 0.3917 | No |
|  | MDMB-CHMICA | MDMB-CHMINACA | -0.0533 | 0.6906 | No | -0.591 | 0.0661 | No |
|  | AM-2201 | THJ-2201 | -0.3743 | 0.0004 | Yes | -0.184 | 0.5046 | No |
| **Overall comparison**<br>(mean Diff ± SEM) | | | **-0.1392**<br>±<br>**0.2225** | **0.5759** | **No** | **0.4733**<br>±**0.1334** | **0.0381** | **Yes** |

**Supplementary Table IV.** SAR analysis for impact of head moiety on hERG efficacy (maximal reduction in Gmax) and potency (IC50). P values for compound-to-compound comparisons are from Extra-sum-of-squares F-test and linear regression. P values for overall comparisons are from two-tailed paired t-tests. Trend denotes overall effects that are not significant, but all go in the same direction.

| | Comparisons | | Efficacy (max reduction in $G_{\max}$) | | | $\log IC_{50}$ | | |
| --- | --- | --- | --- | --- | --- | --- | --- | --- |
|  | Compound 1 | Compound 2 | Mean Diff | P value | Significant? | Mean Diff | P value | Significant? |
| **AB vs. ADB** | AB-FUBINACA | ADB-FUBINACA | -0.2476 | 0.3634 | No | 0.3450 | 0.1409 | No |
|  | AB-CHMINACA | ADB-CHMINACA | -0.1330 | 0.5763 | No | 0.6450 | 0.0505 | No |
|  | AB-CHMICA | ADB-CHMICA | 0.1637 | 0.2726 | No | 0.195 | 0.5921 | No |
| **Overall comparison**<br>(mean Diff ± SEM) | | | **0.0723**<br>±<br>**0.1225** | **0.6150** | **No** | **0.3950**<br>±<br>**0.1323** | **0.0962** | **No**<br>(trend) |
| **AB vs. MDMB** | AB-FUBINACA | MDMB-FUBINACA | -0.2511 | 0.2236 | No | 0.472 | 0.0443 | Yes |
|  | AB-CHMINACA | MDMB-CHMINACA | -0.6017 | 0.0430 | Yes | 0.6040 | 0.0718 | No |
|  | AB-CHMICA | MDMB-CHMICA | 0.0922 | 0.3839 | No | 0.411 | 0.1804 | No |
| **Overall comparison**<br>(mean Diff ± SEM) | | | **-0.2535**<br>±<br>**0.2003** | **0.3331** | **No** | **0.4957**<br>±<br>**0.0567** | **0.0129** | **Yes** |
| **ADB vs. MDMB** | ADB-FUBINACA | MDMB-FUBINACA | -0.0035 | 0.9904 | No | 0.127 | 0.7422 | No |
|  | ADB-CHMINACA | MDMB-CHMINACA | -0.4687 | 0.2146 | No | -0.0410 | 0.9245 | No |
|  | ADB-4en-PINACA | MDMB-4en-PINACA | 0.2509 | 0.1961 | No | -0.2060 | 0.5273 | No |
|  | ADB-5'Br-INACA | MDMB-5'Br-INACA | 0.1302 | 0.6260 | No | -0.0123 | 0.9804 | No |
|  | ADB-5'Br-BUTINACA | MDMB-5'Br-BUTINACA | -0.3093 | 0.1663 | No | 0.808 | 0.0482 | Yes |
|  | ADB-CHMICA | MDMB-CHMICA | -0.0715 | 0.5329 | No | 0.216 | 0.5150 | No |
| **Overall comparison**<br>(mean Diff ± SEM) | | | **-0.0786**<br>±<br>**0.1100** | **0.5067** | **No** | **0.1485**<br>±<br>**0.1446** | **0.3516** | **No** |

**Supplementary Table V.** SAR analysis for impact of core moiety on CB1 receptor efficacy (Emax) and potency (EC50). P values for compound-to-compound comparisons are from Brown-Forsythe and Welch ANOVA tests. P values for overall comparisons are from two-tailed paired t-tests. Trend denotes overall effects that are not significant, but all go in the same direction.

|  | Comparisons |  | Efficacy (E<sub>max</sub>) |  |  | Potency (logEC<sub>50</sub>) |  |  |
| --- | --- | --- | --- | --- | --- | --- | --- | --- |
|  | Compound 1 | Compound 2 | Mean Diff | P value | Significant? | Mean Diff | P value | Significant? |
| **Indole**<br>**vs.**<br>**Indazole** | AB-CHMICA | AB-CHMINACA | 2.081 | 0.9247 | No | 0.3489 | 0.0216 | Yes |
|  | ADB-CHMICA | ADB-CHMINACA | 3.284 | 0.1469 | No | 0.6109 | 0.0058 | Yes |
|  | MDMB-CHMICA | MDMB-CHMINACA | 0.6356 | 0.9998 | No | 0.3405 | 0.0150 | Yes |
|  | AM-2201 | THJ-2201 | 6.900 | 0.0641 | No | -0.2500 | 0.0464 | Yes |
| **Overall comparison**<br>**(mean Diff ± SEM)** |  |  | **-3.225**<br>**±**<br>**1.291** | **0.0879** | **No**<br>**(trend)** | **0.2626**<br>**±**<br>**0.1820** | **0.2448** | **No** |

**Supplementary Table VI.** SAR analysis for impact of head moiety on CB1 receptor efficacy (Emax) and potency (EC50). P values for compound-to-compound comparisons are from Brown-Forsythe and Welch ANOVA tests. P values for overall comparisons are from two-tailed paired t-tests. Trend denotes overall effects that are not significant, but all go in the same direction.

|  | Comparisons |  | Efficacy (E<sub>max</sub>) |  |  | Potency (logEC<sub>50</sub>) |  |  |
| --- | --- | --- | --- | --- | --- | --- | --- | --- |
|  | Compound 1 | Compound 2 | Mean Diff | P value | Significant? | Mean Diff | P value | Significant? |
| **AB**<br>**vs.**<br>**ADB** | AB-FUBINACA | ADB-FUBINACA | 2.638 | 0.8301 | No | 0.2541 | 0.1145 | No |
|  | AB-CHMINACA | ADB-CHMINACA | 1.015 | 0.9991 | No | 0.3663 | 0.0395 | Yes |
|  | AB-CHMICA | ADB-CHMICA | -0.1872 | >0.9999 | No | 0.1043 | 0.2688 | No |
| **Overall comparison**<br>(mean Diff ± SEM) |  |  | **-1.000**<br>**±**<br>**0.5774** | **0.2254** | **No** | **0.2416**<br>**±**<br>**0.07589** | **0.0861** | **No**<br>**(trend)** |
| **AB**<br>**vs.**<br>**MDMB** | AB-FUBINACA | MDMB-FUBINACA | 5.054 | 0.3044 | No | 1.051 | 0.0011 | Yes |
|  | AB-CHMINACA | MDMB-CHMINACA | 1.170 | 0.9983 | No | 0.8071 | 0.0017 | Yes |
|  | AB-CHMICA | MDMB-CHMICA | 2.616 | 0.4704 | No | 0.8155 | 0.0003 | Yes |
| **Overall comparison**<br>(mean Diff ± SEM) |  |  | **3.333**<br>**±**<br>**0.8819** | **0.0634** | **No**<br>**(trend)** | **0.8912**<br>**±**<br>**0.07993** | **0.0079** | **Yes** |
| **ADB**<br>**vs.**<br>**MDMB** | ADB-FUBINACA | MDMB-FUBINACA | 2.416 | 0.8719 | No | 0.7968 | 0.0033 | Yes |
|  | ADB-CHMINACA | MDMB-CHMINACA | 0.1542 | >0.9999 | No | 0.4408 | 0.0120 | Yes |
|  | ADB-4en-PINACA | MDMB-4en-PINACA | 8.179 | 0.1623 | No | 0.0200 | >0.9999 | No |
|  | ADB-5'Br-INACA | MDMB-5'Br-INACA | 1.044 | >0.9999 | No | 0.430 | 0.0049 | Yes |
|  | ADB-5'Br-BUTINACA | MDMB-5'Br-BUTINACA | -3.822 | 0.6589 | No | -0.3675 | 0.0251 | Yes |
|  | ADB-CHMICA | MDMB-CHMICA | 2.803 | 0.2178 | No | 0.7111 | 0.0013 | Yes |
| **Overall comparison**<br>(mean Diff ± SEM) |  |  | **2.000**<br>**±**<br>**1.592** | **0.2644** | **No** | **0.3385**<br>**±**<br>**0.1796** | **0.1181** | **No** |

**Supplementary Table VII.**
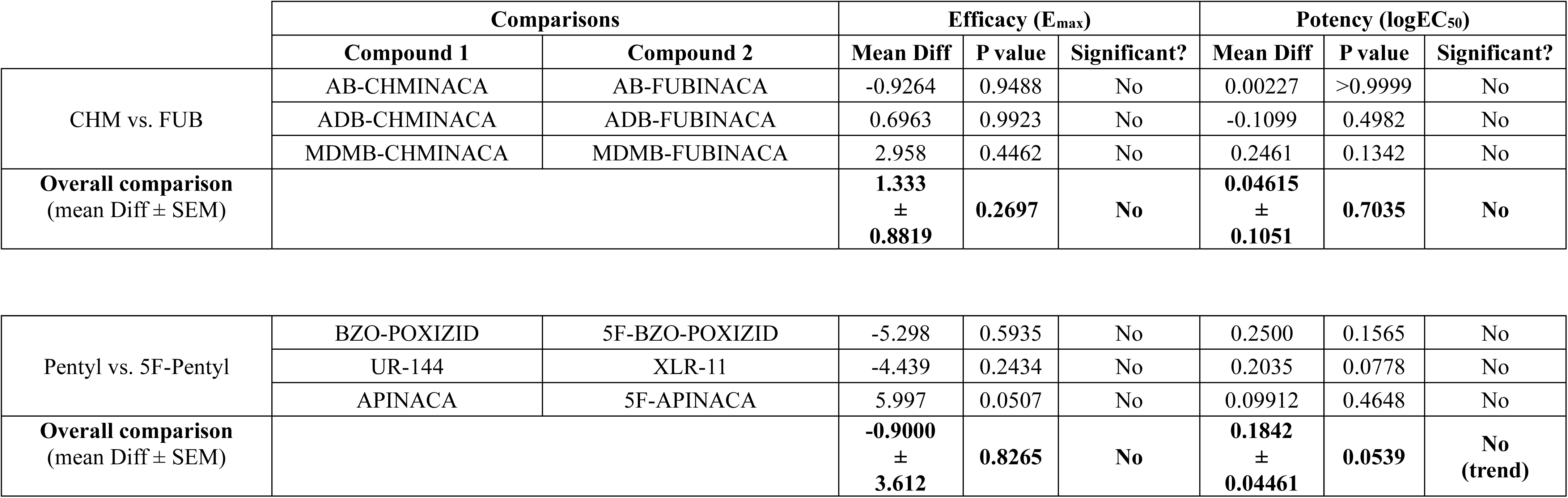
SAR analysis for impact of tail moiety on CB1 receptor efficacy (Emax) and potency (EC50). P values for compound-to-compound comparisons are from Brown-Forsythe and Welch ANOVA tests. P values for overall comparisons are from two-tailed paired t-tests.

## Notes

### Competing Interest Statement

The authors have declared no competing interest.

